# The structure, function, and evolution of a complete human chromosome 8

**DOI:** 10.1101/2020.09.08.285395

**Authors:** Glennis A. Logsdon, Mitchell R. Vollger, PingHsun Hsieh, Yafei Mao, Mikhail A. Liskovykh, Sergey Koren, Sergey Nurk, Ludovica Mercuri, Philip C. Dishuck, Arang Rhie, Leonardo G. de Lima, David Porubsky, Andrey V. Bzikadze, Milinn Kremitzki, Tina A. Graves-Lindsay, Chirag Jain, Kendra Hoekzema, Shwetha C. Murali, Katherine M. Munson, Carl Baker, Melanie Sorensen, Alexandra M. Lewis, Urvashi Surti, Jennifer L. Gerton, Vladimir Larionov, Mario Ventura, Karen H. Miga, Adam M. Phillippy, Evan E. Eichler

**Affiliations:** Department of Genome Sciences, University of Washington School of Medicine, Seattle, WA 98195, USA; Developmental Therapeutics Branch, National Cancer Institute, Bethesda, MD 20892, USA; Genome Informatics Section, Computational and Statistical Genomics Branch, National Human Genome Research Institute, National Institutes of Health, Bethesda, MD 20892, USA; Department of Biology, University of Bari, Aldo Moro, Bari 70121, Italy; Stowers Institute for Medical Research, Kansas City, MO 64110, USA; Graduate Program in Bioinformatics and Systems Biology, University of California, San Diego, CA 92093, USA; McDonnell Genome Institute, Department of Genetics, Washington University School of Medicine, St. Louis, MO 63108, USA; Howard Hughes Medical Institute, University of Washington, Seattle, WA 98195, USA; Department of Pathology, University of Pittsburgh, Pittsburgh, PA 15213, USA; Center for Biomolecular Science and Engineering, University of California, Santa Cruz, Santa Cruz, CA 95064, USA

## Abstract

The complete assembly of each human chromosome is essential for understanding human biology and evolution. Using complementary long-read sequencing technologies, we complete the first linear assembly of a human autosome, chromosome 8. Our assembly resolves the sequence of five previously long-standing gaps, including a 2.08 Mbp centromeric α-satellite array, a 644 kbp defensin copy number polymorphism important for disease risk, and an 863 kbp variable number tandem repeat at chromosome 8q21.2 that can function as a neocentromere. We show that the centromeric α-satellite array is generally methylated except for a 73 kbp hypomethylated region of diverse higher-order α-satellite enriched with CENP-A nucleosomes, consistent with the location of the kinetochore. Using a dual long-read sequencing approach, we complete the assembly of the orthologous chromosome 8 centromeric regions in chimpanzee, orangutan, and macaque for the first time to reconstruct its evolutionary history. Comparative and phylogenetic analyses show that the higher-order α-satellite structure evolved specifically in the great ape ancestor, and the centromeric region evolved with a layered symmetry, with more ancient higher-order repeats located at the periphery adjacent to monomeric α-satellites. We estimate that the mutation rate of centromeric satellite DNA is accelerated at least 2.2-fold, and this acceleration extends beyond the higher-order α-satellite into the flanking sequence.

## INTRODUCTION

Since the announcement of the sequencing of the human genome 20 years ago^1,2^, human chromosomes have remained unfinished due to large regions of highly identical repeats located within centromeres, segmental duplication, and the acrocentric short arms of chromosomes. The presence of large swaths (>100 kbp) of highly identical repeats that are themselves copy number polymorphic has meant that such regions have persisted as gaps, limiting our understanding of human genetic variation and evolution^3,4^. In the case of centromeres, for example, the AT-rich, 171 bp repeat, known as α-satellite, is organized in tandem to form hundreds to thousands of higher-order repeats (HORs) that span mega-base pairs of human DNA and are variable in copy number between homologous chromosomes^5–8^. Such repetitive structures have complicated cloning and assembly of these and other regions of the human genome and, as a result, the sequences have either remained as gaps or are presented as decoys of predicted sequence to improve mapping against the human reference^9,10^.

The advent of long-read sequencing technologies and associated algorithms have now made it possible to systematically assemble these regions from native DNA for the first time^11–13^. In addition, the use of DNA from complete hydatidiform moles (CHMs) to serve as reference genomes has greatly simplified sequence resolution of these complex regions. Most CHMs carry only the paternal complement of human chromosomes due to an aberrant fertilization event in which a single sperm duplicates to give rise to two identical haploid sets of chromosomes. As a result, there is no allelic variation, permitting the assembly of a single haplotype without interference from a second haplotype^14^. The use of long reads from CHM DNA created the first comprehensive map of human structural variation^15^ and the first report of a completely sequenced human X chromosome, where the centromere was fully resolved^16^.

Here, we present the first complete linear assembly of a human autosomal chromosome not only to permit the study of human biology and evolution but to serve as a benchmark for the completion of other chromosomes and future diploid genomes. We chose human chromosome 8 because it carries a modestly sized centromere (approximately 1.5-2.2 Mbp)^8,17^, where the α-satellite repeats are organized into a well-defined HOR array. The chromosome, however, also contains one of the most structurally dynamic regions in the human genome—the β-defensin gene cluster located at 8p23.1^18–20^—as well as a neocentromere located at 8q21.2, which have been largely unresolved for the last 20 years. We use the finished chromosome 8 sequence to perform the first comparative sequence analyses of complete centromeres across the great ape phylogeny and show how this information enables new insights into the structure, function, and evolution of our genome.

## RESULTS

### Telomere-to-telomere assembly of chromosome 8

To resolve the gaps in human chromosome 8 (**Fig. 1a**), we developed a targeted assembly method that leverages the complementary strengths of Oxford Nanopore Technologies (ONT) and Pacific Biosciences (PacBio) long-read sequencing (**Fig. 1b; Methods**). We reasoned that ultra-long (>100 kbp) ONT reads harbor sufficient sequence variation to permit the assembly of complex regions, generating an initial sequence scaffold that could be replaced with highly accurate PacBio high-fidelity (HiFi) contigs to improve the overall base accuracy. To this end, we generated 20-fold sequence coverage of ultra-long ONT data and 32.4-fold coverage of PacBio HiFi data from a CHM (CHM13hTERT; abbr. CHM13; **Extended Data Fig. 1**; **Methods**). Over half of the ultra-long ONT data is composed of reads exceeding 139.8 kbp in length, with the longest mapped read 1.538 Mbp long (**Extended Data Fig. 1a**). More than half of the PacBio HiFi data is contained in reads greater than 17.8 kbp, with a median accuracy exceeding 99.9% (**Extended Data Fig. 1b**). We assembled complex regions in chromosome 8 by first creating a library of singly unique nucleotide k-mers (SUNKs)^21^, or sequences of length *k* that occur approximately once per haploid genome (here, *k* = 20), from CHM13 PacBio HiFi data (**Methods**). These SUNKs were validated with Illumina data generated from the same genome and used to barcode ultra-long ONT reads (**Fig. 1b; Methods**). Ultra-long ONT reads sharing highly similar barcodes were assembled into an initial sequence scaffold that traverses each gap and complex genomic region within chromosome 8 (**Fig. 1b; Methods**). We improved the base-pair accuracy of the sequence scaffolds by replacing the raw ONT sequence with several concordant PacBio HiFi contigs and integrating them into a linear assembly of human chromosome 8 from Nurk and colleagues^11^ (**Fig. 1b; Methods**).

**Figure 1.**
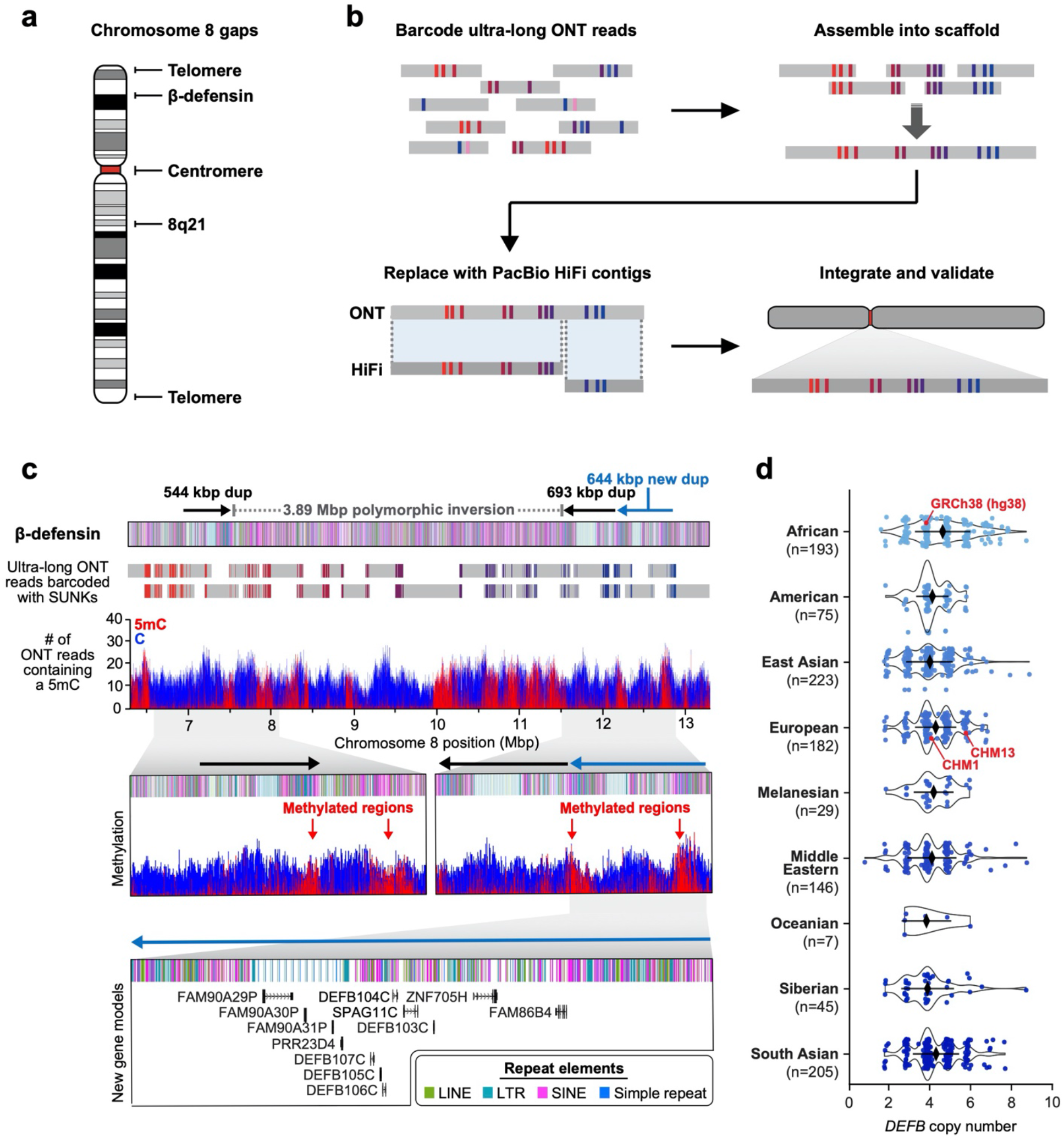
Telomere-to-telomere assembly of human chromosome 8 via a targeted assembly method. **a)** Gaps in the GRCh38 chromosome 8 reference sequence. **b)** Targeted assembly method to resolve complex repeat regions in the human genome. Ultra-long ONT reads (gray) are barcoded with singly unique nucleotide k-mers (SUNKs; colored bars) and assembled into a sequence scaffold. Regions within the scaffold sharing high sequence identity with PacBio HiFi contigs (dark gray) are replaced, thereby improving the base accuracy to >99.99%. The PacBio HiFi assembly is integrated into an assembly of chromosome 8 from the CHM13 genome^11^ and subsequently validated with orthogonal technologies. **c)** Sequence, structure, methylation status, and genetic composition of the CHM13 β-defensin locus. The CHM13 locus contains three segmental duplications (SDs) (dups) located at chr8:7098892-7643091, chr8:11528114-12220905, and chr8:12233870-12878079 in the assembly. A 3,885,023 bp inversion (located at chr8:7643092-11528113) separates the first and second duplication. Although two SDs had been previously reported^20^, the other duplication (light blue) is newly resolved. Resolution of the entire locus was achieved via assembly of 26 ultra-long ONT reads (gray) barcoded with SUNKs (colored bars). The methylation status of the region was determined from mapped ultra-long ONT reads using Nanopolish^25^. 5-methylcytosine (5mC) is indicated in red and unmethylated cytosine is indicated in blue. Iso-Seq data reveal that the new duplication contains twelve new protein-coding genes, five of which are *DEFB* genes (**Extended Data Fig. 15** shows a schematic of all *DEFA*- and *DEFB*-related genes across the β-defensin locus). **d)** Copy number of the *DEFB* genes [chr8:7783837−7929198 in GRCh38 (hg38)] throughout the human population. CHM13 has six copies of *DEFB* genes, one set per SD per haplotype, while CHM1 and GRCh38 only have four copies (red data points).

The complete telomere-to-telomere sequence of human chromosome 8 is 146,259,671 bases long and encompasses 3,334,256 additional bases missing from the current reference genome (GRCh38). Most of the additions reside within distinct chromosomal regions: a ~644 kbp copy number polymorphic β-defensin gene cluster mapping to chromosome 8p23.1 (**Fig. 1c**); the complete centromere corresponding to 2.08 Mbp of α-satellite DNA (**Fig. 2a**); a 863 kbp 8q21.2 variable number tandem repeat (VNTR) (**Fig. 3a**); and both telomeric regions ending with the canonical TTAGGG repeat sequence (**Extended Data Fig. 2**). We validated the organization and accuracy of the chromosome 8 assembly via a suite of orthogonal technologies, including optical mapping (Bionano Genomics), Strand-seq^22,23^, and comparisons to finished BAC sequence as well as whole-genome sequence Illumina data derived from the same source genome (**Methods**). Our analyses show that the CHM13 chromosome 8 assembly is free of assembly errors, false joins, and misorientations (**Extended Data Fig. 3**). We estimate the overall base accuracy to be between 99.9915% and 99.9999% (quality value (QV) score between 40.70 and 63.19, as determined from sequenced BACs and mapped k-mers^24^, respectively). An analysis of 24 million human full-length transcripts generated from Iso-Seq data identifies 61 protein-coding and 33 noncoding loci that map better to this finished chromosome 8 sequence than to GRCh38, including the discovery of novel genes mapping to copy number polymorphic regions (see below; **Fig. 1d**, **Extended Data Fig. 4**).

**Figure 2.**
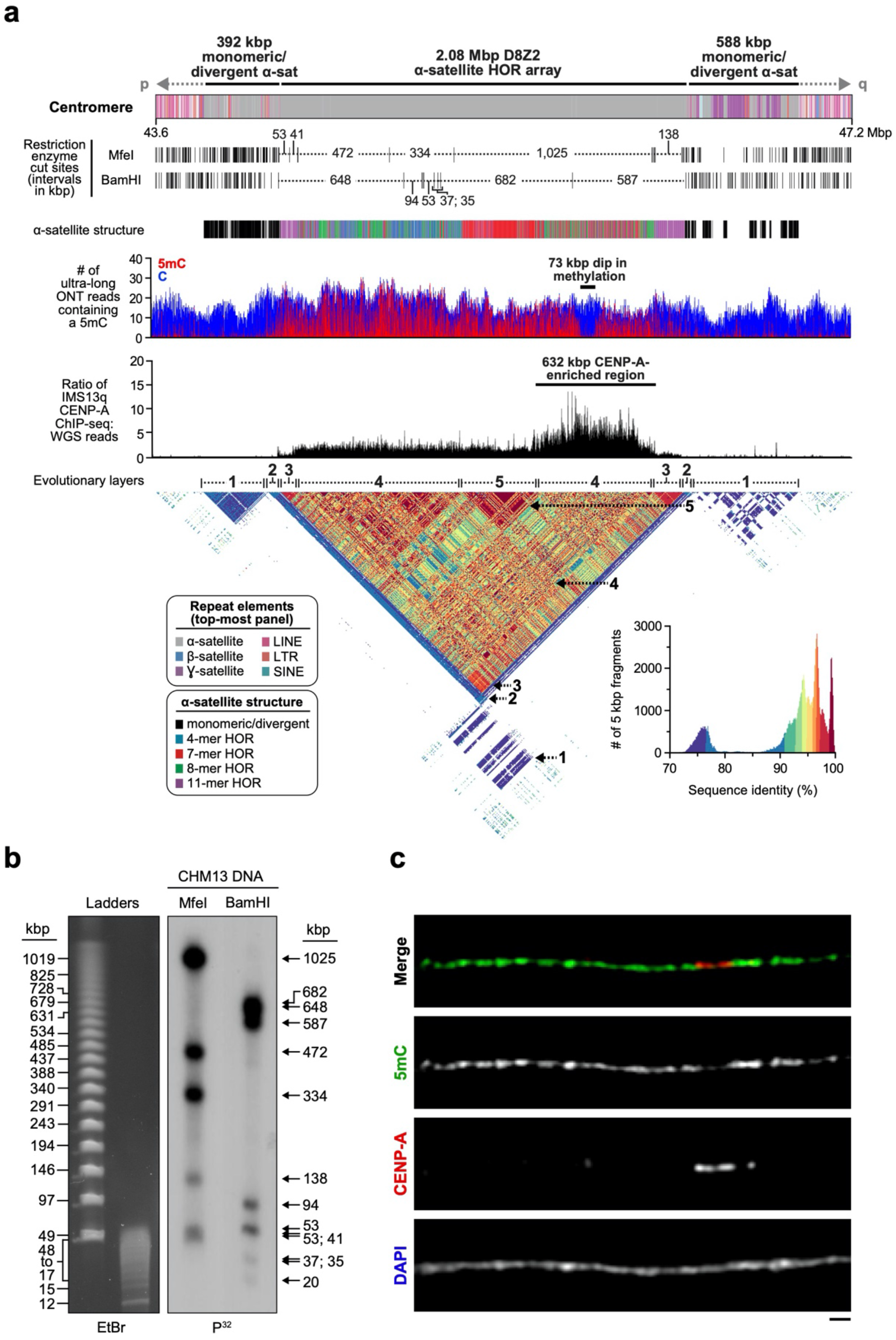
Sequence, structure, and epigenetic map of the chromosome 8 centromeric region. **a)** Schematic showing the composition of the CHM13 chromosome 8 centromere. The centromeric region is comprised of a 2.08 Mbp D8Z2 α-satellite HOR array flanked by regions of monomeric and/or divergent α-satellite interspersed with retrotransposons, β-satellite, and ɣ-satellite. The predicted restriction digest pattern is shown and supported by the pulsed-field gel (PFG) Southern blot in **Panel b**. The D8Z2 α-satellite HOR array is primarily composed of four types of higher-order repeats (HORs; see **Extended Data Fig. 8, Methods** for details) and is heavily methylated except for a 73 kbp hypomethylation region. Mapping of normalized CENP-A ChIP-Seq data from a diploid human genome known as IMS13q^31^ reveals that centromeric chromatin is primarily located within a 632 kbp region encompassing the hypomethylated region (**Extended Data Fig. 9** includes another CENP-A ChIP-seq dataset and details). A pairwise sequence identity map across the centromeric region indicates that the centromere is composed of five distinct evolutionary layers (indicated with dashed arrows). **b)** PFG Southern blot of CHM13 DNA confirms the structure and organization of the chromosome 8 centromeric HOR array indicated in **Panel a**. Left: EtBr staining; Right: P^32^-labeled chromosome 8 α-satellite-specific probe. **c)** Representative images of a CHM13 chromatin fiber showing that CENP-A is enriched in an unmethylated region. Bar = 1 micron.

**Figure 3.**
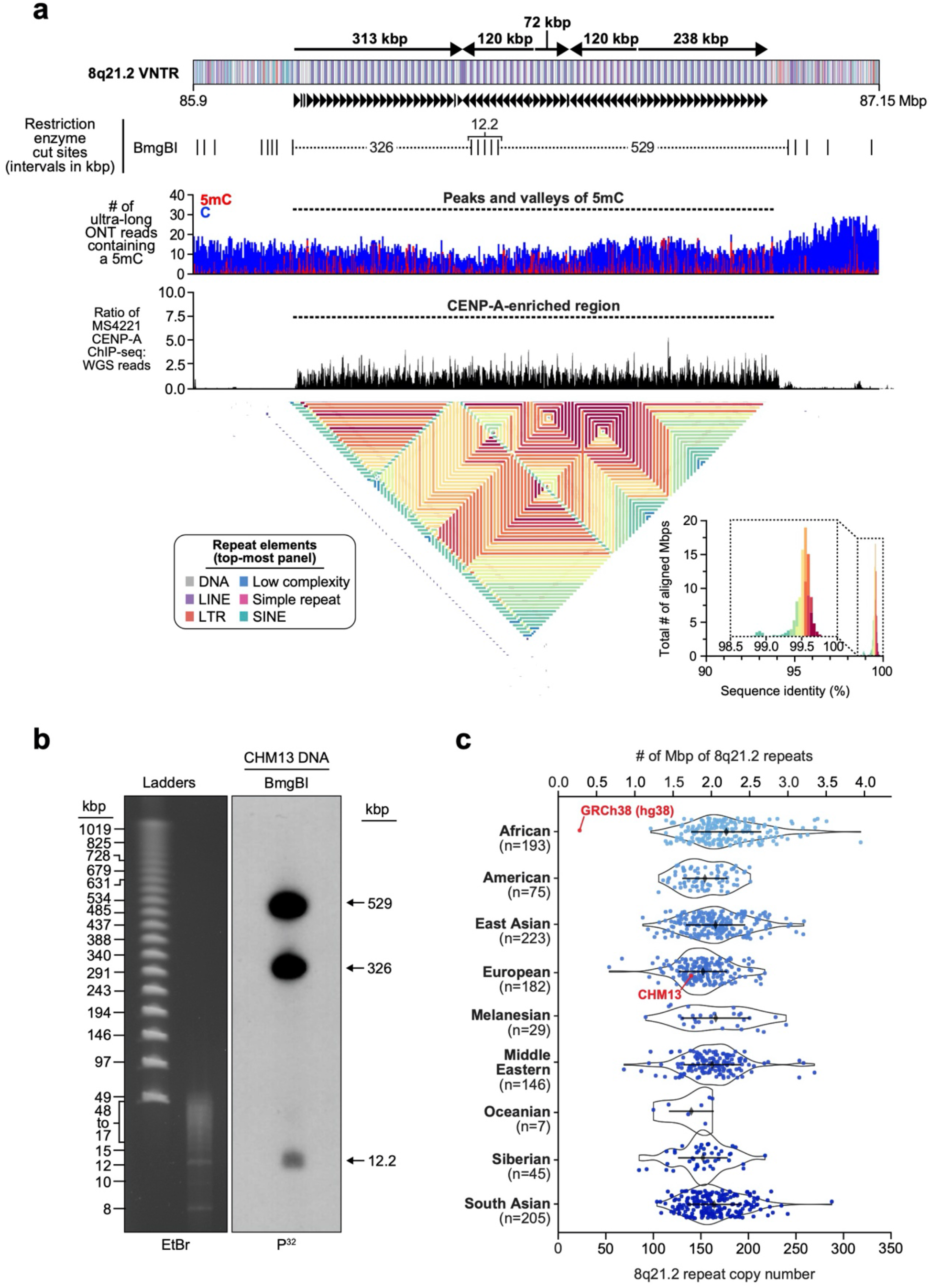
Sequence, structure, and epigenetic map of the neocentromeric chromosome 8q21.2 VNTR. **a)** Schematic showing the composition of the CHM13 8q21.2 VNTR. This VNTR is comprised of 67 full and 7 partial 12.192 kbp repeats that span 863 kbp in total. The predicted restriction digest pattern is indicated. Each repeat is methylated within a 3 kbp region and hypomethylated within the rest of the sequence. Mapping of CENP-A ChIP-seq data from the chromosome 8 neodicentric cell line known as MS4221^31,32^ (**Methods**) reveals that centromeric chromatin is primarily located on the hypomethylated portion of the repeat. A pairwise sequence identity map across the region indicates a mirrored symmetry within a single layer, consistent with the evolutionarily young status of the tandem repeat. **b)** PFG Southern blot of CHM13 DNA digested with BmgBI confirms the size and organization of the chromosome 8q21.2 VNTR. Left: EtBr staining; Right: P^32^-labeled chromosome 8q21.2-specific probe. **c)** Copy number of the 8q21 repeat [chr8:85792897−85805090 in GRCh38] throughout the human population. CHM13 is estimated to have 144 total copies of the 8q21 repeat, or 72 copies per haplotype, while GRCh38 only has 26 copies (red data points).

Our targeted assembly method is particularly useful for traversing large complex regions of highly identical duplications. A case in point is the β-defensin gene cluster^18^, which we resolved into a single 7.06 Mbp locus—substantially larger than the 4.56 Mbp region in the current human reference genome (GRCh38), which is flanked by two 50 kbp gaps (**Fig. 1c**). To assemble this locus, we initially barcoded 26 ultra-long ONT reads (averaging 378.7 kbp in length) with SUNKs and assembled them to generate a 7.06 Mbp sequence scaffold. PacBio HiFi contigs concordant with the ONT-based scaffold replaced 99.9934% of the sequence (7,058,731 out of 7,059,195 bp), increasing the overall base accuracy of the assembly to 99.9911% (QV score of 40.48; estimated with mapped BACs; **Extended Data Fig. 5a**). Our analysis shows that the β-defensin assembly is free from structural errors and misassemblies (**Extended Data Fig. 5a**). Additionally, our analysis reveals a more complex haplotype than GRCh38, consistent with previously published reports of structural variation associated with the chromosome 8p23.1 β-defensin gene cluster^18,20^. We resolve the breakpoints of one of the largest common inversion polymorphisms in the human genome (3.89 Mbp in length) and show that the breakpoints map within large, highly identical duplications that are copy number polymorphic in the human population (**Fig. 1d)**. In contrast to the human reference, which carries two such segmental duplications (SDs), there are three SDs in CHM13: a 544 kbp SD on the distal end and two 693 and 644 kbp SDs on the proximal end, respectively (**Fig. 1c**). Each SD cassette carries at least five β-defensin genes and, as a result, we identify five additional β-defensin genes that are virtually identical at the amino acid level to the reference (**Fig. 1c, Extended Data Table 1)**. Because ONT data allows methylation signals to be assessed^25^, we inferred the methylation status of cytosines across the entire β-defensin locus. All three SDs harbor a 151-163 kbp methylated region residing in the LTR-rich region of the duplication, while the remainder of the SD, including the β-defensin gene cluster, is largely unmethylated (**Fig. 1c**), consistent with its transcription. Complete sequence resolution of this alternate haplotype is important because the inverted haplotype preferentially predisposes to recurrent microdeletions associated with developmental delay, microcephaly, and congenital heart defects^26,27^, and copy number polymorphism of the five β-defensin genes has been associated with immune-related phenotypes such as psoriasis and Crohn’s disease^19,28^.

### Sequence resolution of the chromosome 8 centromere

Prior studies have estimated the length of the chromosome 8 centromere to be between 1.5 and 2.2 Mbp, based on analysis of the HOR α-satellite array^8,17^. While various HORs of different lengths are thought to comprise the centromere, the predominant species has a unit length of 11 monomers, resulting in a tandem repeat of 1881 bp^8,17^. Using our targeted assembly method, we spanned the chromosome 8 centromere with 11 ultra-long ONT reads (mean length 389.4 kbp), which were replaced with PacBio HiFi contigs based on SUNK barcoding. Unlike the ONT assembly, the HiFi assembly was not completely continuous but was more accurate, allowing it to be anchored uniquely into the ONT sequence scaffold. Our assembled CHM13 chromosome 8 centromere consists of a 2.08 Mbp D8Z2 α-satellite HOR array flanked by blocks of monomeric α-satellite on the p- (392 kbp) and q- (588 kbp) arms (**Fig. 2a**). Both monomeric α-satellite blocks are interspersed with LINEs, SINEs, and LTRs, with tracts of ɣ-satellite specific to the q-arm. We validated the sequence, structure, and organization of the chromosome 8 centromere using five orthogonal methods. First, long-read sequence read-depth analysis from two orthogonal native DNA sequencing platforms shows uniform coverage, suggesting that the assembly is free from large structural errors (**Extended Data Fig. 6a**). Fluorescent *in situ* hybridization (FISH) on stretched chromosomes confirms the long-range order and organization of the centromere (**Extended Data Fig. 6a,b**). Droplet digital PCR shows that there are 1344 +/− 142 D8Z2 HORs within the α-satellite array, consistent with our estimates (**Extended Data Fig. 6c; Methods)**. Pulsed-field gel electrophoresis Southern blots on CHM13 DNA digested with two different restriction enzymes recapitulates the banding pattern predicted from the assembly (**Fig. 2a,b**). Finally, applying our assembly approach to ONT and HiFi data available for a diploid human genome generates two additional chromosome 8 centromere haplotypes, replicating the overall organization with only subtle differences in overall length of HOR arrays (**Extended Data Fig. 7**, **Extended Data Table 2**).

Using the assembled centromere sequence, we investigated its genetic and epigenetic organization. On a genetic level, we found that the chromosome 8 centromeric HOR array is primarily composed of four distinct HOR types represented by 4, 7, 8, or 11 α-satellite monomer cassettes (**Fig. 2a, Extended Data Fig. 8**). While the 11-mer predominates (36%), the other HORs are also abundant (19-23%) and are all derivatives of the 11-mer (**Extended Data Fig. 8b,c**). Interestingly, we find that HORs are differentially distributed regionally across the centromere. While most regions are admixed with different HOR types, we also identify regions of homogeneity, such as clusters of 11-mers mapping to the periphery of the HOR array (92 and 158 kbp in length) and a 177 kbp region in the center that is composed solely of 7-mer HORs. To investigate the epigenetic organization, we mapped methylated cytosines along the centromeric region and found that most of the α-satellite HOR array is methylated, except for a small, 73 kbp hypomethylated region (**Fig. 2a**). To determine if this hypomethylated region is the site of the epigenetic centromere (marked by the presence of nucleosomes containing the histone H3 variant, CENP-A), we mapped CENP-A ChIP-seq data from diploid human cells and found that CENP-A was primarily located within a 632 kbp stretch encompassing the hypomethylated region (**Fig. 2a, Extended Data Fig. 9**). Subsequent chromatin fiber FISH revealed that CENP-A maps to the hypomethylated region within the α-satellite HOR array (**Fig. 2c**). Remarkably, the hypomethylated region shows some of the greatest HOR admixture, suggesting a potential optimization of HOR subtypes associated with the active kinetochore (mean entropy over the 73 kbp region = 1.91; **Extended Data Fig. 8a, Methods**).

To better understand the long-range organization and evolution of the centromere, we generated a pairwise sequence identity heat map (**Methods**), which compares the sequence identity of 5 kbp fragments along the length of the centromere (**Fig. 2a**). We find that the centromere consists of five major evolutionary layers that show mirror symmetry. The outermost layer resides in the monomeric α-satellite, where sequences are highly divergent from the rest of the centromere but are more similar to each other (Arrow 1). The second layer defines the monomeric-to-HOR transition and is a short (57-60 kbp) region. The p and q regions are 87-92% identical with each other but only 78% or less with other centromeric satellites (Arrow 2). The third layer is completely composed of HORs. The p and q regions are 92 and 149 kbp in length, respectively, and share more than 96% sequence identity with each other (Arrow 3) but less than that with the rest of the centromere. This layer is composed largely of homogenous 11-mers and defines the transition from unmethylated to methylated DNA. The fourth layer is the largest and defines the bulk of the HOR α-satellite (1.42 Mbp in total). It shows the greatest admixture of different HOR subtypes and, once again, the p and q blocks share identity with each other but are more divergent from the rest of the layers (Arrow 4). Both blocks are highly methylated with the exception of the 73 kbp hypomethylated region mapping to the q-arm. Finally, the fifth layer encompasses the centermost 416 kbp of the HOR array, a region of near-perfect sequence identity that is divergent from the rest of the centromere (Arrow 5).

### Sequence resolution of the chromosome 8q21.2 VNTR

The layered and mirrored nature of the chromosome 8 centromere is reminiscent of another gap region located at chromosome 8q21.2, which we resolved for the first time (**Fig. 3**). This region is a cytogenetically recognizable euchromatic variant^29^ thought to contain one of the largest VNTRs in the human genome^29^. The 12.192 kbp repeating unit encodes the *GOR1/REXO1L1* pseudogene and is highly copy number polymorphic among humans^29,30^. This VNTR is of biological interest because it is the site of a recurrent neocentromere, where a functional centromere devoid of α-satellite has been observed in multiple unrelated individuals^31,32^. The complete genetic and epigenetic composition of the 8q21.2 VNTR has not yet been resolved because the region largely corresponds to a gap in the human reference genome (GRCh38). Using our approach, we successfully assembled the VNTR into an 863.5 kbp sequence composed of ~71 repeating units (67 complete and 7 partial units) (**Fig. 3a**). A pulsed-field gel Southern blot of digested CHM13 DNA confirms the length and structure of the VNTR (**Fig. 3a,b**). Chromatin fiber FISH estimates that the array is composed of 67 +/− 5.2 repeats, consistent with the assembly (**Extended Data Fig. 10, Methods**). Mapping of long-read data reveals uniform coverage along the entire assembly (**Extended Data Fig. 10a**), indicating a lack of large structural errors. We estimate that the 12.192 kbp repeat unit varies from 53 to 158 copies in the human population, creating tandem repeat arrays ranging from 652 kbp to 1.94 Mbp (**Fig. 3c**). We identify a higher-order structure of the VNTR consisting of five distinct domains that alternate in orientation (**Fig. 3a**), and each domain contains 5 to 23 complete repeat units that are more than 98.5% identical to each other (**Fig. 3a**). Mapping of methylated cytosines to the array shows that each 12.192 kbp repeat is primarily methylated in the 3 kbp region corresponding to *GOR1*/*REXO1L1*, while the rest of the repeat unit is largely unmethylated (**Fig. 3a**). Mapping of centromeric chromatin from a cell line harboring an 8q21.2 neocentromere^32^ shows that the CENP-A nucleosomes map to the unmethylated region of the repeat unit in the CHM13 assembly (**Fig. 3a**). While this is consistent with the VNTR being the potential site of the functional kinetochore of the neocentromere, sequence and assembly of the neocentromere-containing cell line will be critically important.

### Centromere evolutionary reconstruction

We used the complete sequence of the centromeric region of chromosome 8 to comparatively target the orthologous regions in other primate species in an effort to fully reconstruct the evolutionary history of the centromere over the last 25 million years. We first assembled reference genomes corresponding to chimpanzee, orangutan, and macaque genomes. Each diploid genome assembly was sequenced to 25- to 40-fold coverage using PacBio HiFi sequence data, with which we generated assemblies ranging from 6.02 to 6.12 Gbp in size, consistent with assemblies where both haplotypes were assembled (**Extended Data Table 3**). Focusing on the centromere, we also generated ONT datasets for the same references, which were simultaneously used to construct an initial sequence scaffold of each orthologous region corresponding to the human chromosome 8 centromere. Once the scaffold assembly was established and barcoded with SUNKs, we used these SUNKs to replace the ONT scaffold with overlapping high-accuracy PacBio HiFi contigs. We successfully generated two contiguous assemblies of the chimpanzee chromosome 8 centromere (one for each haplotype), one haplotype assembly from the orangutan chromosome 8 centromere, and one complete haplotype from the macaque chromosome 8 centromere (**Fig. 4**). Mapping of long-read data to each assembly shows uniform coverage, indicating a lack of large structural errors (**Extended Data Figs. 11,12)**. Analysis of each nonhuman primate (NHP) chromosome 8 centromere reveals distinct HOR patterns ranging in size from 1.69 Mbp in chimpanzee to 10.92 Mbp in macaque, consistent with estimates from short-read sequence data and cytogenetic analyses^33,34^ (**Fig. 4**).

**Figure 4.**
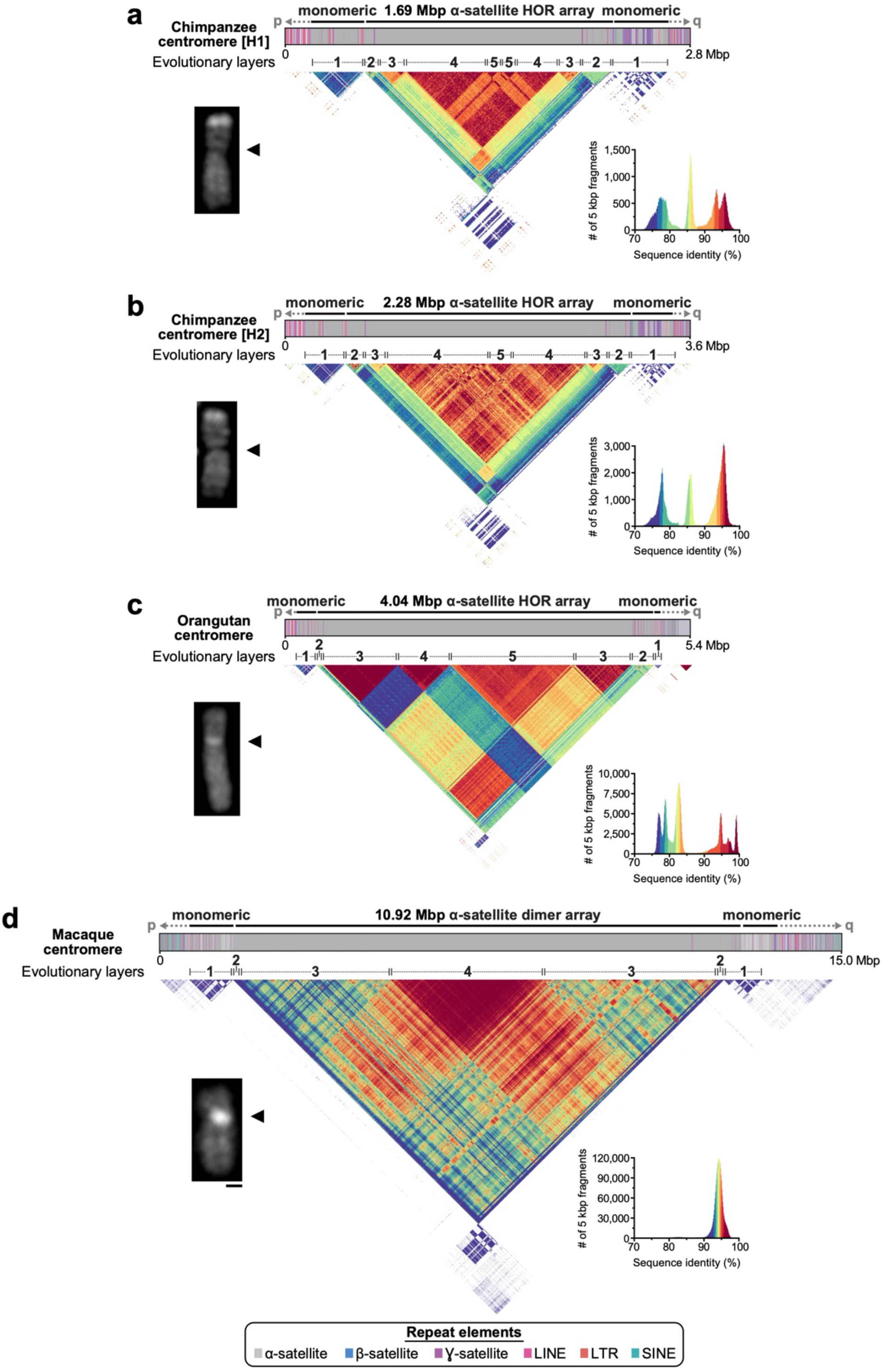
Sequence and structure of the chimpanzee, orangutan, and macaque chromosome 8 centromeres. **a-d)** Structure and sequence identity of the chimpanzee H1 (**Panel a**), chimpanzee H2 (**Panel b**), orangutan (**Panel c**), and macaque (**Panel d**) chromosome 8 centromeres. Each centromere has a mirrored organization consisting of either four or five distinct evolutionary layers. The size of each centromeric region is consistent with microscopic analyses, showing increasingly bright DAPI staining with increasing centromere size. Bar = 1 micron.

Similar to human, we constructed a pairwise sequence identity map of each NHP centromere. The data, once again, reveal a mirrored and layered organization, with the chimpanzee organization being most similar to human (**Figs. 2a, 4**). In general, each NHP chromosome 8 centromere is composed of four or five distinct layers, with the outermost layer showing the lowest degree of sequence identity (73-78% in chimpanzee and orangutan; 90-92% in macaque) and the innermost layer showing the highest sequence identity (90-100% in chimpanzee and orangutan; 94-100% in macaque). The orangutan structure is striking in that there appears to be very little admixture of HOR units between the layers, in contrast to other apes where the different HOR cassettes are derived from a major HOR structure. The blocks of orangutan HORs (with the exception of layer 3) show a reduced degree of sequence identity. This suggests that the orangutan centromere evolved as a mosaic of independent HOR units. In contrast to all apes, the macaque lacks HORs and, instead, harbors a basic dimeric repeat structure^33^, which is much more homogenous and highly identical (>90%) across the nearly 11 Mbp of assembled centromeric array.

We assessed the phylogenetic relationship between higher-order and monomeric α-satellites from each primate centromere using a maximum-likelihood framework, taking advantage of the positional information from the completed sequence to define orthologous locations between the species (**Fig. 5a**). We find that all great ape higher-order α-satellite sequences (corresponding to layers 2-5) cluster into a single clade, while the monomeric α-satellite (layer 1) split into two clades separated by tens of millions of years. The proximal clade contains monomeric α-satellite from both the p- and q-arms, while the more divergent clade shares monomeric α-satellite solely from the q-arm, and specifically, the α-satellite nestled between clusters of ɣ-satellite (**Extended Data Fig. 13**). Unlike great apes, both monomeric and dimeric repeat structures from the macaque group together and are sister clades to the monomeric ape clades, suggesting a common ancient origin restricted to these flanking pericentromeric regions.

**Figure 5.**
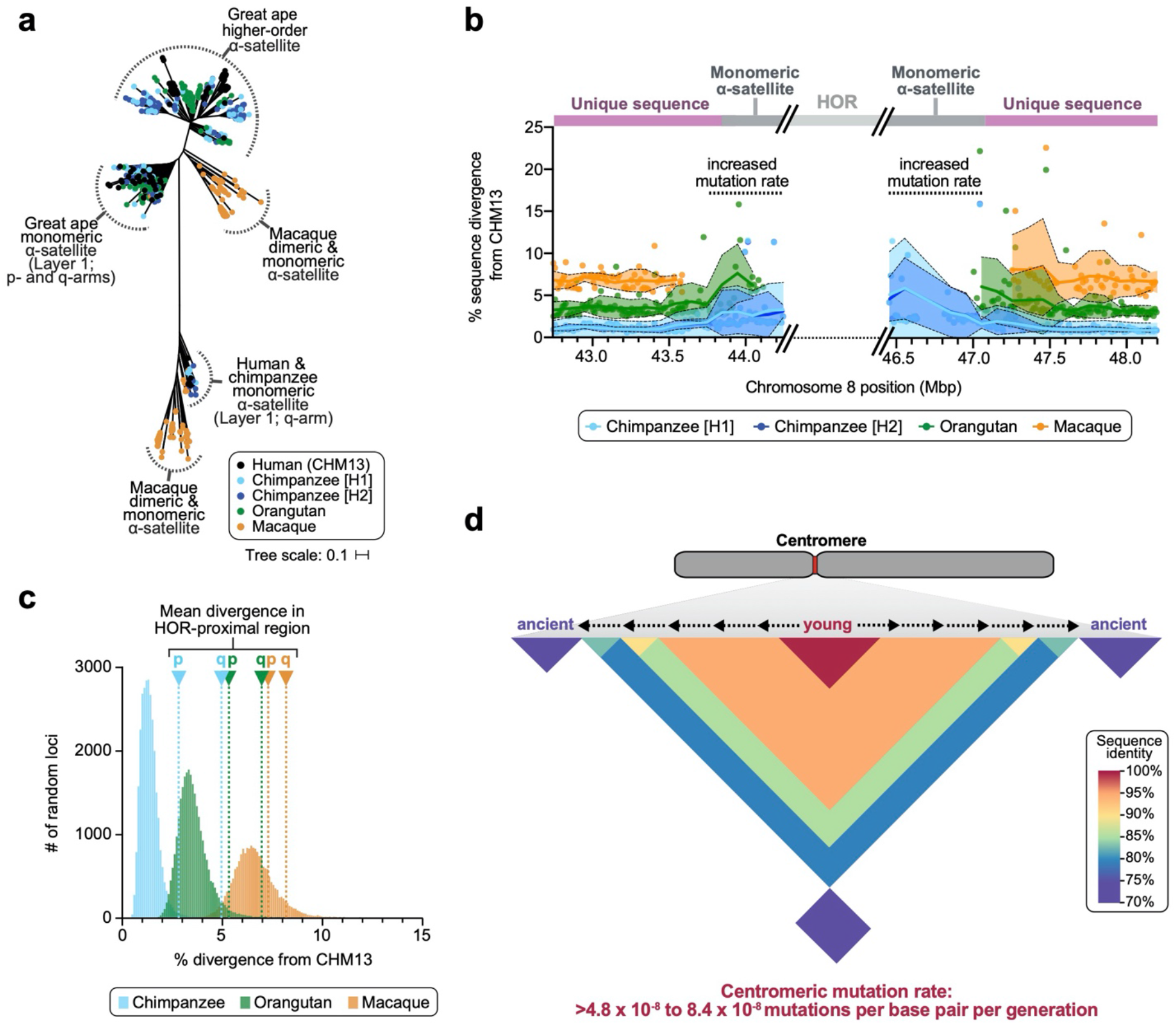
Evolution of the chromosome 8 centromere. **a)** Phylogenetic tree of human, chimpanzee, orangutan, and macaque α-satellites from the HOR and monomeric regions of the chromosome 8 centromere. A portion of the human and chimpanzee monomeric α-satellite is evolutionarily closer to the macaque α-satellite (bottom of the tree; see **Extended Data Fig. 13** for bootstrapping annotations). **b)** Plot showing the sequence divergence between the CHM13 and nonhuman primates (NHPs) in the regions flanking the chromosome 8 α-satellite HOR array. The mean and standard deviation (bold line and shaded region) are calculated over a sliding window of 200 kbp with a 100 kbp overlap. Individual data points from 10 kbp pairwise sequence alignments are shown. **c)** Histogram of the sequence divergence between CHM13 and chimpanzee, orangutan, or macaque at thousands of random 10 kbp loci. **d)** Model of centromere evolution. Centromeric α-satellite HORs evolve in the center of the array via unequal crossing over and homogenization, pushing older, more ancient HORs to the edges, consistent with hypotheses previously put forth^38,39,41^. The centromeric mutation rate is estimated to be at least 4.8 to 8.4 × 10^−8^ mutations per base pair per generation, which is 2.2 to 3.8 higher than the mean mutation rate measured from nearly 20,000 random loci.

Because we independently assembled the centromere for each primate and successfully transitioned from α-satellite to unique sequence for both the p- and q-arms, we used this orthology to understand how rapidly sequences decay over the course of evolution. Anchored in orthologous sequence, we assessed divergence based on 10 kbp windows of pairwise alignments in the ~2 Mbp flanking the α-satellite HOR array (**Fig. 5b**). We find that the mean divergence increases more than threefold as the sequence transitions from unique to monomeric α-satellite. Such increases are rare in the genome based on sampling of at least 19,926 random loci, where only 1.27-1.99% of loci show comparable levels of divergence (**Fig. 5c**). Using evolutionary models (**Methods**), we estimate a minimal mutation rate of the chromosome 8 centromeric region of ~4.8 × 10^−8^ and ~8.4 × 10^−8^ mutations per base pair per generation on the p- and q-arms, respectively, which is 2.2- to 3.8-fold higher than the basal mean mutation rate (~2.2 × 10^−8^) (**Extended Data Table 4**). These analyses provide the first complete comparative sequence analysis of a primate centromere for an orthologous chromosome and a framework for future studies of genetic variation and evolution of these regions across the genome.

## DISCUSSION

Chromosome 8 is the first human autosome to be sequenced and assembled from telomere to telomere and contains only the third completed human centromere to date^16,35^. The assembly of chromosome 8 was achieved via a dual-technology assembly method that leverages the scaffolding capability of ONT with the high-accuracy of PacBio HiFi long reads to traverse complex regions of our genome that have remained as gaps since human genomes were first assembled^1,2^. The result is a whole-chromosome assembly with an estimated base-pair accuracy exceeding 99.99%. We also successfully applied this hybrid strategy to reconstruct centromeric regions from diploid organisms. We generated complete draft assemblies of both chromosome 8 centromere haplotypes, for example, from a chimpanzee and a human sample. In contrast, only one haplotype was contiguously assembled in macaque and orangutan with the second haplotype remaining incomplete. It should be noted that both the human and chimpanzee diploid samples are genetically admixed, and it is possible that this heterogeneity facilitated the partitioning of reads and the reconstruction of both haplotypes from these samples.

Comparison of the centromeric regions between human chromosomes 8 and X reveals similarities and differences in their epigenetic status and organization^16^. Both chromosomes harbor a pocket of hypomethylation (~61-73 kbp in length), and we show that this hypomethylated region is enriched for the centromeric histone, CENP-A—although CENP-A enrichment extends over a broader swath (632 kbp) with its peak centered over the hypomethylated region. These data strongly suggest we have identified the functional kinetochore binding site^36,37^. In contrast to the X chromosome HOR array, which is primarily comprised of one type of HOR^16^, the chromosome 8 centromere shows a mixture of primarily four different types of HORs that are present in near-equal abundance and organized into layers of differing sequence identity. Although the HOR units are derived from the original 11-mer repeat, the degree of admixture and purity varies considerably across the centromere, suggesting a more complex model of evolution. While this layered HOR organization is evident for both chromosomes X and 8, the mirror symmetry is only observed for chromosome 8. Other centromeres will need to be sequenced and assembled to determine the generality of this feature (**Extended Data Fig. 14**). Importantly, the X chromosome HOR array is less than half as diverse when compared to chromosome 8^16^, likely due to the slower rate of mutation of the X chromosome compared to human autosomes.

The layered and mirrored organization of the centromere is consistent with rapid evolutionary turnover of centromeric repeats^38–40^, wherein highly identical repeats undergo unequal crossover and homogenization, pushing older, more divergent repeats to the edges in an assembly-line fashion (**Fig. 5d**). The chromosome 8 centromere reveals five such layers, with the evolutionarily youngest layer in the center of the α-satellite HOR array and more ancient layers flanking it. The two additional human chromosome 8 centromeres, as well as each primate centromere, show a similar gradient of divergence as one proceeds towards the periphery, with some of the most identical tracts mapping centrally. The location and purity of the 7-mer HOR units in the human chromosome 8 centromeric array, for example, are consistent with the Smith model of rapid unequal crossing over and homogenization. Surprisingly, the hypomethylated regions, which we predict define the active kinetochore, do not map to the most active site of homogenization as postulated by the library hypothesis^41^. In contrast, the 73 kbp hypomethylated region maps to a segment showing some of the greatest admixture, suggesting that these different HOR subtypes may be important for defining the functional centromere in contrast to the most identical HOR tract. The “mosaic” architecture of the orangutan HOR may be the result of a recent or even ongoing arms race to define the most competent HOR associated with the kinetochore in that species. The assembled sequence allows such hypotheses to be functionally tested in the future.

In addition to the centromere, we resolved other complex loci involving copy number variable SDs and VNTRs. The new copies are predicted to encode novel duplicate genes (e.g., new copies of defensin genes) and, as such, the additional sequence adds to our understanding of human gene annotation and, subsequently, access to the underlying genetic variation therein. The completion of these sequences further enhances functional annotation of the genome, showing, for example, that the 12.192 kbp tandem repeat defines the site of kinetochore attachment in individuals with a chromosome 8 neocentromere. In addition, analysis of the sequence structure provides potential insights into the ontogeny of centromeres. For instance, the complete sequence of this 863 kbp VNTR shows that it possesses a higher-order repetitive structure where adjacent segments flip between a direct and indirect orientation in blocks ranging from 72 to 313 kbp units (**Fig. 3a)**. Several studies of neocentromeres devoid of satellite sequences have suggested that such inversions are commonly shared features among some ectopic centromeres^42,43^, thus providing further support that such inverted configurations may be a key feature for new centromere formation.

Our targeted assembly method also facilitated draft assemblies of centromeres and their heterochromatic flanking sequences from closely related NHPs. This allowed orthologous relationships to be established and phylogenetic relationships among satellite repeats to be determined with respect to their location. We confirm that HOR structures evolved after apes diverged from Old World monkeys (OWM; <25 million years ago)^33,44,45^ but also distinguish different classes of monomeric repeats that share an ancient origin with the OWM. One ape monomeric clade present only in the q-arm clearly groups with the macaque’s (**Extended Data Fig. 13**). We hypothesize that this ~70 kbp segment present in chimpanzee and human, but absent in orangutan, represents the remnants of the ancestral centromere of apes and OWM now residing only at the pericentromeric periphery of chromosome 8q. The other shared monomeric clade (mapping to layer 1) continues to have diverged after apes split from OWM and likely represents the origin of ape HORs. This observation supports the emergence of a new class of monomers in great apes with greater potential to form centromeres^40^. In contrast to apes, OWM show a much simpler trajectory of continual decay of a basic dimeric HOR^46^ from a 4 Mbp core of near-perfect sequence identity, creating a satellite more than double in size when compared to ape counterparts.

Using orthologous sequence alignments for the heterochromatin transition regions, we estimate that mutation rates increase by two to fourfold in proximity to the HOR, likely due to the action of concerted evolution, unequal crossing-over, and saltatory amplification^33,39,40^. This acceleration includes monomeric satellites with some evidence that it extends beyond the satellites themselves up to ~30 kbp and ~170 kbp into unique regions on the p- and q-arms, respectively. It should be noted that our current centromere mutation rate estimates represent a lower bound for the overall centromere, as they are calculated from the monomeric α-satellite rather than the HOR array itself. Even between the human and chimpanzee lineage separated by six million years, the α-satellite HOR array is too divergent to generate a simple pairwise alignment that would permit the computation of a mutation rate over these sequences. Novel satellite repeats are homogenized and swept to fixation within a population through mechanisms such as unequal crossing over and gene conversion^47^. Rapid evolution of centromeres has been described in multiple species as a driving force for speciation due to the accumulation of sequence differences that result in highly divergent HOR sequences, and subsequently, causes incompatibility and reproductive barrier in hybrids between closely related species^48–50^. In this light, it is interesting that sequence comparisons among three human centromere 8 haplotypes predict regions of excess allelic variation and structural divergence (**Extended Data Fig. 7c-e**), although the locations within the HOR differ among haplotypes (**Extended Data Fig. 7**). Now that complex regions such as these can be sequenced and assembled, it will be important to extend these analyses to other centromeres, multiple individuals, and additional species to understand their full impact with respect to genetic variation and evolution.

## METHODS

### Cell culture

CHM13hTERT (abbr. CHM13) cells were cultured in complete AmnioMax C-100 Basal Medium (Thermo Fisher Scientific, 17001082) supplemented with 15% AmnioMax C-100 Supplement (Thermo Fisher Scientific, 12556015) and 1% penicillin-streptomycin (Thermo Fisher Scientific, 15140122). Chimpanzee (*Pan troglodytes*; Clint; S006007) and macaque (*Macaque mulatta*; AG07107) cells were cultured in MEM α containing ribonucleosides, deoxyribonucleosides, and L-glutamine (Thermo Fisher Scientific, 12571063) supplemented with 12% FBS (Thermo Fisher Scientific, 16000-044) and 1% penicillin-streptomycin (Thermo Fisher Scientific, 15140122). Orangutan (*Pongo abelii*; Susie; PR01109) cells were cultured in MEM α containing ribonucleosides, deoxyribonucleosides, and L-glutamine (Thermo Fisher Scientific, 12571063) supplemented with 15% FBS (Thermo Fisher Scientific, 16000-044) and 1% penicillin-streptomycin (Thermo Fisher Scientific, 15140122). All cells were cultured in a humidity-controlled environment at 37°C with 95% O_2_ and 5% CO_2_.

### DNA extraction, library preparation, and sequencing

PacBio HiFi data were generated from the chimpanzee, orangutan, and macaque genomes as previously described^51^ with modifications. Briefly, high-molecular-weight (HMW) DNA was extracted from cells using a modified Qiagen Gentra Puregene Cell Kit protocol^52^. HMW DNA was used to generate HiFi libraries via the SMRTbell Express Template Prep Kit v2 and SMRTbell Enzyme Clean Up kits (PacBio). Size selection was performed with SageELF (Sage Science), and fractions sized 11, 14, 18, 22, or 25 kbp (as determined by FEMTO Pulse (Agilent)) were chosen for sequencing. Libraries were sequenced on the Sequel II platform with three to seven SMRT Cells 8M (PacBio) using either Sequel II Sequencing Chemistry 1.0 and 12-hour pre-extension or Sequel II Sequencing Chemistry 2.0 and 3- or 4-hour pre-extension, both with 30-hour movies, aiming for a minimum estimated coverage of 25X in HiFi reads (assuming a genome size of 3.2 Gbp). Raw data was processed using the CCS algorithm (v3.4.1 or v4.0.0) with the following parameters: ‒minPasses 3 ‒minPredictedAccuracy 0.99 ‒maxLength 21000 or 50000.

Ultra-long ONT data were generated from the CHM13, chimpanzee, and orangutan genomes according to a previously published protocol^53^. Briefly, 5 × 10^7^ cells were lysed in a buffer containing 10 mM Tris-Cl (pH 8.0), 0.1 M EDTA (pH 8.0), 0.5% w/v SDS, and 20 ug/mL RNase A for 1 hour at 37C. Proteinase K (200 ug/mL) was added, and the solution was incubated at 50C for 2 hours. DNA was purified via two rounds of 25:24:1 phenol-chloroform-isoamyl alcohol extraction followed by ethanol precipitation. Precipitated DNA was solubilized in 10 mM Tris (pH 8) containing 0.02% Triton X-100 at 4C for two days. Libraries were constructed using the Rapid Sequencing Kit (SQK-RAD004) from ONT with modifications to the manufacturer’s protocol. Specifically, 2-3 ug of DNA was resuspended in a total volume of 18 ul with 16.6% FRA buffer. FRA enzyme was diluted 2- to 12-fold into FRA buffer, and 1.5 uL of diluted FRA was added to the DNA solution. The DNA solution was incubated at 30C for 1.5 min, followed by 8C for 1 min to inactivate the enzyme. RAP enzyme was diluted 2- to 12-fold into RAP buffer, and 0.5 uL of diluted RAP was added to the DNA solution. The DNA solution was incubated at room temperature (RT) for 2 hours before loading onto a primed FLO-MIN106 R9.4.1 flow cell for sequencing.

Additional ONT data was generated from the CHM13, chimpanzee, orangutan, and macaque genomes. Briefly, HMW DNA was extracted from cells using a modified Qiagen Gentra Puregene Cell Kit protocol^52^. HMW DNA was prepared into libraries with the Ligation Sequencing Kit (SQK-LSK109) from ONT and loaded onto primed FLO-MIN106 or FLO-PRO002 R9.4.1 flow cells for sequencing on the GridION or PromethION, respectively. All ONT data were base called with Guppy 3.6.0 or 4.11.0 with the HAC model.

### PacBio HiFi whole-genome assembly

Chimpanzee, orangutan, and macaque genomes were assembled from PacBio HiFi data (**Extended Data Table 2**) using HiCanu^11^ (v2.0). The CHM13 genome was previously assembled with HiCanu and described by Nurk and colleagues^11^. Contigs from each primate assembly were used to replace the ONT-based sequence scaffolds in targeted regions (described below).

### Targeted sequence assembly

Gapped regions within human chromosome 8 were targeted for assembly via a SUNK-based method that combines both PacBio HiFi and ONT data. Specifically, CHM13 PacBio HiFi data was used to generate a library of SUNKs (*k = 20*; total = 2,062,629,432) via Jellyfish (v2.2.4) based on the sequencing coverage of the HiFi dataset. 99.88% (2,060,229,331) of the CHM13 PacBio HiFi SUNKs were validated with CHM13 Illumina data (SRR3189741). A subset of CHM13 ultra-long ONT reads aligning to the CHM1 β-defensin patch (GenBank: KZ208915.1) or select regions within the GRCh38 chromosome 8 reference sequence (chr8:42,881,543-47,029,467 for the centromere and chr8:85,562,829-85,848,463 for the 8q21.2 locus) were barcoded with Illumina-validated SUNKs. Reads sharing at least 50 SUNKs were selected for inspection to determine if their SUNK barcodes overlapped. SUNK barcodes can be composed of “valid” and “invalid” SUNKs. Valid SUNKs are those that occur once in the genome and are located at the exact position on the read. In contrast, invalid SUNKs are those that occur once in the genome but are falsely located at the position on the read, and this may be due to a sequencing or base-calling error, for example. Valid SUNKs were identified within the barcode as those that share pairwise distances with at least ten other SUNKs on the same read. Reads that shared a SUNK barcode containing at least three valid SUNKs and their corresponding pairwise distances (+/−1% of the read length) were assembled into a tile. The process was repeated using the tile and subsetted ultra-long ONT reads several times until a sequence scaffold spanning the gapped region was generated. Validation of the scaffold organization was carried out via three independent methods. First, the sequence scaffold and underlying ONT reads were subjected to RepeatMasker (v3.3.0) to ensure that read overlaps were concordant in repeat structure. Second, the centromeric scaffold and underlying ONT reads were subjected to StringDecomposer^54^ to validate the HOR organization in overlapping reads. Finally, the sequence scaffold for each target region was incorporated into the CHM13 chromosome 8 assembly^11^, thereby filling the gaps in the chromosome 8 assembly. CHM13 PacBio HiFi and ONT data were aligned to the entire chromosome 8 assembly via pbmm2 (v1.1.0) (for PacBio data; https://github.com/PacificBiosciences/pbmm2) or Winnowmap^55^ (v1.0) (for ONT data) to identify large collapses or misassemblies. Although the ONT-based scaffolds are structurally accurate, they are only 87-98% accurate at the base level due to base-calling errors in the raw ONT reads^13^. Therefore, we sought to improve the base accuracy of the sequence scaffolds by replacing the ONT sequences with PacBio HiFi contigs assembled from the CHM13 genome^11^, which have a consensus accuracy greater than 99.99%^11^. Therefore, we aligned CHM13 PacBio HiFi contigs generated via HiCanu^11^ to the chromosome 8 assembly via minimap2^56^ (v2.17-r941; parameters: minimap2 -t 8 -I 8G -a --eqx -x asm20 -s 5000) to identify contigs that share high sequence identity with the ONT-based sequence scaffolds. A typical scaffold had multiple PacBio HiFi contigs that aligned to regions within it. Therefore, the scaffold was used to order and orient the PacBio HiFi contigs and bridge gaps between them when necessary. PacBio HiFi contigs with high sequence identity replaced almost all regions of the ONT-based scaffolds: ultimately, the chromosome 8 assembly is comprised of 146,254,195 bp of PacBio HiFi contigs and only 5,490 bp of ONT sequence scaffolds (99.9963% PacBio HiFi contigs and 0.0037% ONT scaffold). The chromosome 8 assembly was incorporated into a whole-genome assembly of CHM13^11^ for validation via orthogonal methods (detailed below). The chimpanzee, orangutan, and macaque chromosome 8 centromeres were assembled via the same SUNK-based method.

### Accuracy estimation

The accuracy of the CHM13 chromosome 8 assembly was estimated from mapped k-mers using Merqury^24^. Briefly, Merqury (v1.1) was run on the chromosome 8 assembly with the following command: eval/qv.sh CHM13.k21.meryl chr8.fasta chr8_v9.

CHM13 Illumina data (SRR1997411, SRR3189741, SRR3189742, SRR3189743) was used to identify k-mers with *k* = 21. In Merqury, every k-mer in the assembly is evaluated for its presence in the Illumina k-mer database, with any k-mer missing in the Illumina set counted as base-level ‘error’. We detected 1,474 k-mers found only in the assembly out of 146,259,650, resulting in a QV score of 63.19, estimated as follows:

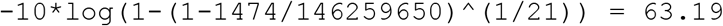

The accuracy percentage for chromosome 8 was estimated from this QV score as:

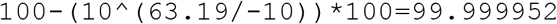

The accuracy of the CHM13 chromosome 8 assembly and β-defensin locus were also estimated from sequenced BACs. Briefly, 66 BACs from the CHM13 chromosome 8 (BAC library VMRC59) were aligned to the chromosome 8 assembly via minimap2^56^ (v2.17-r941) with the following parameters: -I 8G −2K 1500m --secondary=no -a --eqx -Y -x asm20 -s 200000 -z 10000,1000 -r 50000 -O 5,56 -E 4,1 -B 5. QV was then estimated using the CIGAR string in the resulting BAM, counting alignment differences as errors according to the following formula:

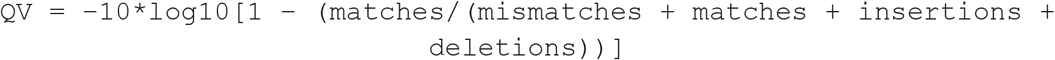

The median QV was 40.6988 for the entire chromosome 8 assembly and 40.4769 for the β-defensin locus (chr8:6300000-13300000; estimated from 47 individual BACs; see **Extended Data Fig. 4** for more details), which falls within the 95% confidence interval for the whole chromosome This QV score was used to estimate the base accuracy^51^ as follows:

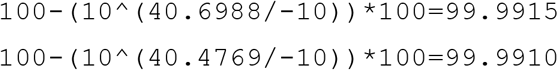

The BAC QV estimation should be considered a lower bound, since differences between the BACs and the assembly may originate from errors in the BAC sequences themselves. Vollger and colleagues showed that BACs can occasionally contain sequencing errors that are not supported by the underlying PacBio HiFi reads^51^. Additionally, the upper bound for the estimated BAC QV is limited to ~53, since BACs are typically ≲200 kbp and, as a result, the maximum calculable QV is 1 error in 200 kbp (QV 53). We also note that the QV of the centromeric region could not be estimated from BACs due to biases in BAC library preparation, which preclude centromeric sequences in BAC clones.

### Strand-seq analysis

We evaluated the directional and structural contiguity of CHM13 chromosome 8 assembly, including the centromere, using Strand-seq data. First, all Strand-seq libraries produced from the CHM13 genome^51^ were aligned to the CHM13 assembly, including chromosome 8 using BWA-MEM^57^ (v0.7.17-r1188) with default parameters for paired-end mapping. Next, duplicate reads were marked by sambamba^58^ (v0.6.8) and removed before subsequent analyses. We used SAMtools^59^ (v1.9) to sort and index the final BAM file for each Strand-seq library. To detect putative misassembly breakpoints in the chromosome 8 assembly, we ran breakpointR^60^ on all BAM files to detect strand-state breakpoints. Misassemblies are visible as recurrent changes in strand state across multiple Strand-seq libraries^61^. To increase our sensitivity of misassembly detection, we created a ‘composite file’ that groups directional reads across all available Strand-seq libraries^62,63^. Next, we ran breakpointR on the ‘composite reads file’ using the function ‘runBreakpointr’ to detect regions that are homozygous (‘ww’; ‘HOM’ – all reads mapped in minus orientation) or heterozygous inverted (‘wc’, ‘HET’ – approximately equal number of reads mapped in minus and plus orientation). To further detect any putative chimerism in the chromosome 8 assembly, we applied Strand-seq to assign 200 kbp long chunks of the chromosome 8 assembly to unique groups corresponding to individual chromosomal homologues using SaaRclust^61,64^. For this, we used the SaaRclust function ‘scaffoldDenovoAssembly’ on all BAM files.

### Bionano analysis

Bionano Genomics data was generated from the CHM13 genome^16^. Long DNA molecules labeled with Bionano’s Direct Labeling Enzyme were collected on a Bionano Saphyr Instrument to a coverage of 130X. The molecules were assembled with the Bionano assembly pipeline Solve3.4, using the nonhaplotype-aware parameters and GRCh38 as the reference. The resulting data produced 261 genome maps with a total length of 2921.6 Mbp and a genome map N50 of 69.02 Mbp.

The molecule set and the nonhaplotype-aware map were aligned to the CHM13 draft assembly and the GRCh38 assembly, and discrepancies were identified between the Bionano maps and the sequence references using scripts in the Bionano software package -- runCharacterize.py, runSV.py, and align_bnx_to_cmap.py.

A second version of the map was assembled using the haplotype-aware parameters. This map was also aligned to GRCh38 and the final CHM13 assembly to verify heterozygous locations. These regions were then examined further.

Analysis of Bionano alignments revealed three heterozygous sites within chromosome 8 located at approximately chr8:21,025,201, chr8:80,044,843, and chr8:121,388,618 (**Extended Data Table 5**). The structure with the greatest ONT read support was selected for inclusion in the chromosome 8 assembly (**Extended Data Table 5**).

### TandemMapper and TandemQUAST analysis of the centromeric HOR array

We assessed the structure of the CHM13 and NHP centromeric HOR arrays by applying TandemMapper and TandemQUAST^65^, which can detect large structural assembly errors in repeat arrays. For the CHM13 centromere, we first aligned ONT reads longer than 50 kbp to the CHM13 assembly containing the contiguous chromosome 8 with Winnowmap^55^ (v1.0) and extracted reads aligning to the centromeric HOR repeat array (chr8:44243868-46323885). We then inputted these reads in the following TandemQUAST command: tandemquast.py -t 24 --nano {ont_reads.fa} -o {out_dir} chr8.fa. For the NHP centromeres, we aligned ONT reads to the whole-genome assemblies containing the contiguous chromosome 8 centromeres with Winnowmap^55^ (v1.0) and extracted reads aligning to the centromeric HOR repeat arrays. We then inputted these reads in the following TandemQUAST command: tandemquast.py -t 24 --nano {ont_reads.fa} -o {out_dir} chr8.fa.

### Methylation analysis

Nanopolish^25^ (v0.12.5) was used to measure CpG methylation from raw ONT reads (>50 kbp in length for CHM13) aligned to whole-genome assemblies via Winnowmap^55^ (v1.0). Nanopolish distinguishes 5-methylcytosine from unmethylated cytosine via a Hidden Markov Model (HMM) on the raw nanopore current signal. The methylation caller generates a log-likelihood value for the ratio of probability of methylated to unmethylated CpGs at a specific k-mer. We filtered methylation calls using the nanopore_methylation_utilities tool (https://github.com/isaclee/nanopore-methylation-utilities)^66^, which uses a log-likelihood ratio of 2.5 as a threshold for calling methylation. CpG sites with log-likelihood ratios greater than 2.5 (methylated) or less than −2.5 (unmethylated) are considered high quality and included in the analysis. Reads that do not have any high-quality CpG sites are filtered from the BAM for subsequent methylation analysis. Nanopore_methylation_utilities integrates methylation information into the BAM file for viewing in IGV’s^67^ bisulfite mode, which was used to visualize CpG methylation.

### Iso-Seq data generation and sequence analyses

RNA was purified from approximately 1 × 10^7^ CHM13 cells using an RNeasy kit (Qiagen; 74104) and prepared into Iso-Seq libraries following a standard protocol^68^. Libraries were loaded on two SMRT Cells 8M and sequenced on the Sequel II. The data were processed via isoseq3 (v8.0), ultimately generating 3,576,198 full-length non-chimeric (FLNC) reads. Poly-A trimmed transcripts were aligned to this CHM13 chr8 assembly and to GRCh38 with minimap2^56^ (v2.17-r941) with the following parameters: -ax splice -f 1000 --sam-hit-only --secondary=no --eqx. Transcripts were assigned to genes using featureCounts^69^ with GENCODE^70^ (v34) annotations, supplemented with CHESS v2.2^71^ for any transcripts unannotated in GENCODE. Each transcript was scored for percent identity of its alignment to each assembly, requiring 90% of the length of each transcript to align to the assembly for it to count as aligned. For each gene, non-CHM13 transcripts’ percent identity to GRCh38 was compared to the CHM13 chromosome 8 assembly. Genes with an improved representation in the CHM13 assembly were identified with a cutoff of 20 improved reads per gene, with at least 0.2% average improvement in percent identity. GENCODE (v34) transcripts were lifted over to the CHM13 chr8 assembly using Liftoff^72^ to compare the GRCh38 annotations to this assembly and Iso-Seq transcripts.

We combined the 3.6 million full-length transcript data (above) with 20,937,742 FLNC publicly available human Iso-Seq data (**Extended Data Table 6**). In total, we compared the alignment of 24,513,940 FLNC reads from 13 tissue and cell line sources to both the completed CHM13 chromosome 8 assembly and the current human reference genome, GRCh38. Of the 848,048 non-CHM13 cell line transcripts that align to chromosome 8, 93,495 (11.02%) align with at least 0.1% greater percent identity to the CHM13 assembly, and 52,821 (6.23%) to GRCh38. This metric suggests that the chromosome 8 reference improves human gene annotation by ~4.79% even though most of those changes are subtle in nature. Overall, 61 protein-coding and 33 noncoding loci have improved alignments to the CHM13 assembly compared to GRCh38, with >0.2% average percent identity improvement, and at least 20 supporting transcripts (**Extended Data Fig. 4, Extended Data Table 7**). As an example, *WDYHV1* (*NTAQ1*) has four amino acid replacements, with 13 transcripts sharing the identical open reading frame to CHM13 (**Extended Data Fig. 16**).

### Pairwise sequence identity heat maps

To generate pairwise sequence identity heat maps, we fragmented the centromere assemblies into 5 kbp fragments (e.g., 1-5000, 5001-10000, etc.) and made all possible pairwise alignments between the fragments using the following minimap2^56^ (v2.17-r941) command: minimap2 -f 0.0001 -t 32 -X --eqx -ax ava-ont. The sequence identity was determined from the CIGAR string of the alignments and then visualized using ggplot2 (geom_raster) in R (v1.1.383)^73^. The color of each segment was determined by sorting the data by identity and then creating 10 equally-sized bins, each of which received a distinct color from the spectral pallet. The choice of a 5 kbp window came after testing a variety of window sizes. Ultimately, we found 5 kbp to be a good balance between resolution of the figure (since each 5 kbp fragment is plotted as a pixel) and sensitivity of minimap2 (fragments less than 5 kbp often missed alignments with the ava-ont preset).

### Analysis of α-satellite organization

To determine the organization of the chromosome 8 centromeric region, we employed two independent approaches. First, we subjected the CHM13 centromere assembly to an *in silico* restriction enzyme digestion wherein a set of restriction enzyme recognition sites were identified within the assembly. In agreement with previous findings that XbaI digestion can generate a pattern of HORs within the chromosome 8 HOR array^17^, we found that each α-satellite HOR could be extracted via XbaI digestion. The *in silico* digestion analysis indicates that the chromosome 8 centromeric HOR array is comprised of 1462 HOR units: 283 4-mers, 4 5-mers, 13 6-mers, 356 7-mers, 295 8-mers, and 511 11-mers. As an alternative approach, we subjected the centromere assembly to StringDecomposer^54^ using a set of 11 α-satellite monomers derived from a chromosome 8 11-mer HOR unit. The sequence of the α-satellite monomers used are as follows: A: AGCATTCTCAGAAACACCTTCGTGATGTTTGCAATCAAGTCACAGAGTTGAACCTTCCGTTTCATAGAGCAGGTTGGAAACACTCTTATTGTAGTATCTGGAAGTGGACATTTGGAGCGCTTTCAGGCCTATGGTGAAAAAGGAAATATCTTCCCATAAAAACGACATAGA; B: AGCTATCTCAGGAACTTGTTTATGATGCATCTAATCAACTAACAGTGTTGAACCTTTGTACTGACAGAGCACTTTGAAACACTCTTTTTTGGAATCTGCAAGTGGATATTTGGATCGCTTTGAGGATTTCGTTGGAAACGGGATGCAATATAAAACGTACACAGC; C: AGCATACTCAGAAAATACTTTGCCATATTTCCATTCAAGTCACAGAGTGGAACATTCCCATTCATAGAGCAGGTTGGAAACACTCTTTTTGGAGTATCTGGAAGTGGACATTTGGAGCGCTTTCTGAACTATGGTGAAAAAGGAAATATCTTCCAATGAAAACAAGACAGA; D: AGCATTCTGAGAAACTTATTTGTGATGTGTGTCCTCAACAAACGGACTTGAACCTTTCGTTTCATGCAGTACTTCTGGAACACTCTTTTTGAAGATTCTGCATGCGGATATTTGGATAGCTTTGAGGATTTCGTTGGAAACGGGCTTACATGTAAAAATTAGACAGC; E: AGCATTCTCAGAAACTTCTTTGTGGTGTCTGCATTCAAGTCACAGAATTGAACTTCTCCTCACATAGAGCAGTTGTGCAGCACTCTATTTGTAGTATCTGGAAGTGGACATTTGGAGGGCTTTGTAGCCTATCTGGAAAAAGGAAATATCTTCCCATGAATGCGAGATAGA; F: AGTAATCTCAGAAACATGTTTATGCTGTATCTACTCAACTAACTGTGCTGAACATTTCTATTGATAGAGCAGTTTTGAGACCCTCTTCTTTTGGAATCTGCAAGTGGATATTTGGATAGATTTGAGGATTTCGTTGGAAACGGGATTATATATAAAAAGTAGACAGC; G: AGCATTCTCAGAAACTTCTTTGTGATGTTTGCATCCAGCTCTCAGAGTTGAACATTCCCTTTCATAGAGTAGGTTTGAAACCCTCTTTTTATAGTGTCTGGAAGCGGGCATTTGGAGCGCTTTCAGGCCTATGCTGAAAAAGGAAATATCTACATATAGAAACTAGACAGA; H: AGCATTCTGAGAATCAAGTTTGTGATGTGGGTACTCAACTAACAGTGTTGATCCATTCTTTTGATACAGCAGTTTTGAACCACACTTTTTGTAGAATCTGCAAGTGGATATTTGGATAGCTGTGAGGATTTCGTTGGAAACGGGAATGTCTTCATAGAAAATTTAGACAGA; I: AGCATTCTCAGAACCTTGATTGTGATGTGTGTTCTCCACTAACAGAGTTGAACCTTTCTTTTGACAGAACTGTTCTGAAACATTCTTTTTATAGAATCTGGAAGTGGATATTTGGAAAGCTTTGAGGATTTCGTTGGAAACGGGAATATCTTCAAATAAAATCTAGCCAGA; J: AGCATTCTAAGAAACATCTTAGGGATGTTTACATTCAAGTCACAGAGTTGAACATTCCCTTTCACAGAGCAGGTTTGAAACAATCTTCTCGTACTATCTGGCAGTGGACATTTTGAGCTCTTTGGGGCCTATGCTGAAAAAGGAAATATCTTCCGACAAAAACTAGTCAGA; K: AGCATTCGCAGAATCCCGTTTGTGATGTGTGCACTCAACTGTCAGAATTGAACCTTGGTTTGGAGAGAGCACTTTTGAAACACACTTTTTGTAGAATCTGCAGGTGGATATTTGGCTAGCTTTGAGGATTTCGTTGGAAACGGTAATGTCTTCAAAGAAAATCTAGACAGA.

This analysis indicated that the chromosome 8 centromeric HOR array is comprised of 1512 HOR units: 283 4-mers, 12 6-mers, 366 7-mers, 303 8-mers, 3 10-mers, 539 11-mers, 2 12-mers, 2 13-mers, 1 17-mer, and 1 18-mer, which is concordant with the *in silico* restriction enzyme digestion results. The predominant HOR types from StringDecomposer are presented in **Extended Data Fig. 8**.

### Copy number estimation

To estimate the copy number for the 8q21.2 VNTR and *DEFB* loci in human lineages, we applied a read-depth based copy number genotyper^21^ to a collection of 1,112 published high-coverage genomes^74–79^. Briefly, sequencing reads were divided into multiples of 36-mer, which were then mapped to a repeat-masked human reference genome (GRCh38) using mrsFAST^80^ (v3.4.1). To increase the mapping sensitivity, we allowed up to two mismatches per 36-mer. The read depth of mappable sequences across the genome was corrected for underlying GC content, and copy number estimate for the locus of interest was computed by summarizing over all mappable bases for each sample.

### Entropy calculation

To define regions of increased admixture within the centromeric HOR array, we calculated the entropy using the frequencies of the different HOR units in 10-unit windows (1 unit slide) over the entire array. The formula for entropy is:

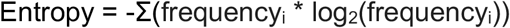

where frequency is (# of HORs) / (total # of HORs) in a 10-unit window. The analysis is analogous to that performed by Gymrek and colleagues^81^.

### Droplet digital PCR

Droplet digital PCR was performed on CHM13 genomic DNA to estimate the number of D8Z2 α-satellite HORs, as was previously done for the DXZ1 α-satellite HORs^16^. Briefly, genomic DNA was isolated from CHM13 cells using the DNeasy Blood & Tissue Kit (Qiagen). DNA was quantified using a Qubit Fluorometer and the Qubit dsDNA HS Assay (Invitrogen). 20 uL reactions were prepared with 0.1 ng of gDNA for the D8Z2 assay or 1 ng of gDNA for the *MTUS1* single-copy gene (as a control). EvaGreen droplet digital PCR (Bio-Rad) master mixes were simultaneously prepared for the D8Z2 and *MTUS1* reactions, which were then incubated for 15 minutes to allow for restriction digest, according to the manufacturer’s protocol.

### Pulsed-field gel electrophoresis and Southern blot

CHM13 genomic DNA was prepared in agarose plugs and digested with either BamHI or MfeI (to characterize the chromosome 8 centromeric region) or BmgBI (to characterize the chromosome 8q21.2 region) in the buffer recommended by the manufacturer. The digested DNA was separated with the CHEF Mapper system (Bio-Rad; autoprogram, 5-850 kbp range, 16 hr run), transferred to a membrane (Amersham Hybond-N+) and blot-hybridized with a 156 bp probe specific to the chromosome 8 centromeric α-satellite or 8q21.2 region. The probe was labeled with P^32^ by PCR-amplifying a synthetic DNA template #233: 5’- TTTGTGGAAGTGGACATTTCGCTTTGTAGCCTATCTGGAAAAAGGAAATATCTTCCCATGAATGCGAGATAGAAGTAATCTCAGAAACATGTTTATGCTGTATCTACTCAACTAACTGTGCTGAACATTTCTATTGTAAAAATAGACAGAAGCATT-3’ (for the centromere of chromosome 8); #264: 5’- TTTGTGGAAGTGGACATTTCGCCCGAGGGGCCGCGGCAGGGATTCCGGGGGACCGGGAGTGGGGGGTTGGGGTTACTCTTGGCTTTTTGCCCTCTCCTGCCGCCGGCTGCTCCAGTTTCTTTCGCTTTGCGGCGAGGTGGTAAAAATAGACAGAAGCATT-3’ (for the organization of the chromosome 8q21.2 locus) with PCR primers #129: 5’-TTTGTGGAAGTGGACATTTC-3’ and #130: 5’- AATGCTTCTGTCTATTTTTA-3’. The blot was incubated for 2 hr at 65°C for pre-hybridization in Church’s buffer (0.5 M Na-phosphate buffer containing 7% SDS and 100 μg/ml of unlabeled salmon sperm carrier DNA). The labeled probe was heat denatured in a boiling water bath for 5 min and snap-cooled on ice. The probe was added to the hybridization Church’s buffer and allowed to hybridize for 48 hr at 65°C. The blot was washed twice in 2× SSC (300 mM NaCl, 30 mM sodium citrate, pH 7.0), 0.05% SDS for 10 min at room temperature, twice in 2× SSC, 0.05% SDS for 5 min at 60°C, twice in 0.5× SSC, 0.05% SDS for 5 min at 60°C, and twice in 0.25× SSC, 0.05% SDS for 5 min at 60°C. The blot was exposed to X-ray film for 16 hr at −80°C.

### Immunofluorescence (IF) and fluorescence *in situ* hybridization (FISH) on chromatin fibers

To determine the location of CENP-A relative to methylated DNA (specifically, 5-methylcytosines), we performed IF on stretched CHM13 chromatin fibers as previously described with modifications^82,83^. Briefly, CHM13 cells were swollen in a hypotonic buffer consisting of a 1:1:1 ratio of 75 mM KCl, 0.8% NaCitrate, and dH_2_O for 5 min. 3.5 × 10^4^ cells were cytospun onto an ethanol-washed glass slide at 800 rpm for 4 min with high acceleration and allowed to adhere for 1 min before immersing in a salt-detergent-urea lysis buffer (25 mM Tris pH 7.5, 0.5 M NaCl, 1% Triton X-100, and 0.3 M urea) for 15 min at room temperature. The slide was slowly removed from the lysis buffer over a time period of 38 s and subsequently washed in PBS, incubated in 4% formaldehyde in PBS for 10 min, and washed with PBS and 0.1% Triton X-100. The slide was rinsed in PBS and 0.05% Tween-20 (PBST) for 3 min, blocked for 30 min with IF block (2% FBS, 2% BSA, 0.1% Tween-20, and 0.02% NaN_2_), and then incubated with a mouse monoclonal anti-CENP-A antibody (1:200, Enzo, ADI-KAM-CC006-E) and rabbit monoclonal anti-5-methylcytosine antibody (1:200, RevMAb, RM231) for 3 h at room temperature. Cells were washed 3x for 5 min each in PBST and then incubated with Alexa Fluor 488 goat anti-rabbit (1:200, Thermo Fisher Scientific, A-11034) and Alexa Fluor 594 conjugated to goat anti-mouse (1:200, Thermo Fisher Scientific, A-11005) for 1.5 h. Cells were washed 3× for 5 min each in PBST, fixed for 10 min in 4% formaldehyde, and washed 3x for 1 min each in dH2O before mounting in Vectashield containing 5 μg/ml DAPI. Slides were imaged on an inverted fluorescence microscope (Leica DMI6000) equipped with a charge-coupled device camera (Leica DFC365 FX) and a 40× 1.4 NA objective lens.

To assess the repeat organization of the 8q21 neocentromere, we performed FISH^84^ on CHM13 chromatin fibers. DNA fibers were obtained following Henry H. Q. Heng’s protocol with minor modifications^85^. Briefly, chromosomes were fixed with methanol:acetic acid (3:1) and dropped on previously clean slides. Chromosomes were dropped onto slides and soaked in PBS 1x. Manual elongation was performed by coverslip and NaOH:ethanol (5:2) solution. We quantified the number and intensity of the probe signals on a set of CHM13 chromatin fibers using ImageJ’s Gel Analysis tool (v1.51) and found that there were 63 +/− 7.55 green signals and 67 +/− 5.20 red signals (n=3 independent experiments), consistent with the 67 full and 7 partial repeats in the CHM13 8q21.2 VNTR.

To validate the organization of the chromosome 8 centromere, we performed FISH on CHM13 cytospun metaphase chromosome spreads in order to increase the chromosome length and improve the resolution of the experiments. We followed the Haaf and Ward protocol^86^ with slight modifications. Briefly, cells were treated with colcemid and resuspended in HCM buffer (10 mM HEPES pH7.3, 30 mM glycerol, 1 mM CaCl_2_, 0.8 mM MgCl_2_) and after 10 minutes, cytospun on silanized slides. Incubation overnight in cold methanol was required to fully fix the chromosomes.

The probes used in the FISH experiments were picked from the human large-insert clone fosmid library ABC10. ABC10 end sequences were mapped using MEGABLAST (similarity=0.99, parameters: -D 2 -v 7 -b 7 -e 1e-40 -p 80 -s 90 -W 12 -t 21 -F F) to a repeat-masked CHM13 genome assembly containing the complete chromosome 8 (parameters: -e wublast -xsmall -no_is -s -species Homo sapiens).

Expected insert size for fosmids was set to (min) 32 kbp and (max) 48 kbp. Resulting clone alignments were grouped into the following categories based on uniqueness of the alignment for a given pair of clones, alignment orientation and the inferred insert size from the assembly.

1. Concordant best: unique alignment for clone pair, insert size within expected fosmid range, expected orientation
2. Concordant tied: non-unique alignment for clone pair, insert size within expected fosmid range, expected orientation
3. Discordant best: unique alignment of clone pair, insert size too small, too large or in opposite expected orientation of expected fosmid clone
4. Discordant tied: non unique alignment for clone pair, insert size too small, too large or in opposite expected orientation of expected fosmid clone
5. Discordant trans: clone pair has ends mapping to different contigs

Clones aligning to regions within the chromosome 8 centromeric region or 8q21.2 locus were selected for FISH validation. The fosmid clones used for validation of the chromosome 8 centromeric region are: 174552_ABC10_2_1_000046302400_C7 for the monomeric α-satellite region, 171417_ABC10_2_1_000045531400_M19 for the entire D8Z2 HOR array, 174222_ABC10_2_1_000044375100_H13 for the central portion of the D8Z2 HOR array. The clones used for validation of the 8q21.2 locus are: 174552_ABC10_2_1_000044787700_O7 for Probe 1 and 173650_ABC10_2_1_000044086000_F24 for Probe 2.

### CENP-A ChIP-seq analysis

We mapped previously published CENP-A ChIP-seq and whole-genome sequencing (WGS) data using two different approaches: 1) BWA-MEM^87^, and 2) a k-mer-based mapping approach we developed (described below). Both results were highly concordant, as shown in **Extended Data Fig. 9**. Diploid datasets used in this analysis include MS4221 CENP-A ChIP-seq and WGS data (SRX246078, SRX246081) and IMS13q CENP-A ChIP-seq and WGS data (SRX246077, SRX246080).

For BWA-MEM mapping, CENP-A ChIP-seq and WGS data were aligned to the CHM13 whole-genome assembly^11^ containing the contiguous chromosome 8 with the following parameters: bwa mem -k 50 -c 1000000 {index} {read1.fastq.gz}for single-end data, and bwa mem -k 50 -c 1000000 {index} {read1.fastq.gz} {read2.fastq.gz} for paired-end data. The resulting SAM files were filtered using SAMtools^59^ with FLAG score 2308 to prevent multi-mapping of reads. With this filter, reads mapping to more than one location are randomly assigned a single mapping location, thereby preventing mapping biases in highly identical regions. The ChIP-seq data were normalized with deepTools^88^ bamCompare with the following parameters: bamCompare -b1 {ChIP.bam} -b2 {WGS.bam} --operation ratio --binSize 1000 -o {out.bw}. The resulting bigWig file was visualized on the UCSC Genome Browser using the CHM13 chromosome 8 assembly as an assembly hub.

For the k-mer-based mapping, the initial BWA-MEM alignment was used to identify reads specific to the chromosome 8 centromeric region (chr8:43600000-47200000). K-mers (*k* = 50) were identified from each chromosome 8 centromere-specific dataset using Jellyfish (v2.3.0) and mapped back onto reads and chromosome 8 centromere assembly allowing for no mismatches. Approximately 93-98% of all k-mers identified in the reads were also found within the D8Z2 HOR array. Each k-mer from the read data was then placed once at random between all sites in the HOR array that had a perfect match to that k-mer. These data were then visualized using a histogram with a bin width of 500 in R (R core team, 2020).

### Phylogenetic analysis

To assess the phylogenetic relationship between α-satellite repeats, we first masked every non-α-satellite repeat in the human and NHP centromere assemblies using RepeatMasker^89^ (v4.1.0). Then, we subjected the masked assemblies to StringDecomposer^54^ using a set of 11 α-satellite monomers derived from a chromosome 8 11-mer HOR unit (described in the “Analysis of α-satellite organization subsection” above). This tool identifies the location of α-satellite monomers in the assemblies, and we used this to extract the α-satellite monomers from the HOR/dimeric array and monomeric regions into multi-FASTA files. We ultimately extracted 12,989, 8,132, 12,224, 25,334, and 63,527 α-satellite monomers from the HOR/dimeric array in human, chimpanzee (H1), chimpanzee (H2), orangutan, and macaque, respectively, and 2,879, 3,781, 3,351, 1,573, and 8,127 monomers from the monomeric regions in human, chimpanzee (H1), chimpanzee (H2), orangutan and macaque, respectively. We randomly selected 100 and 50 α-satellite monomers from the HOR/dimeric array and monomeric regions and aligned them with MAFFT^90,91^ (v7.453). We used IQ-TREE^92^ to reconstruct the maximum-likelihood phylogeny with model selection and 1000 bootstraps. The resulting tree file was visualized in iTOL^93^.

To estimate sequence divergence along the pericentromeric regions, we first mapped each NHP centromere assembly to the CHM13 centromere assembly using minimap2^56^ (v2.17-r941) with the following parameters: -ax asm20 --eqx -Y -t 8 -r 500000. Then, we generated a BED file of 10 kbp windows located within the CHM13 centromere assembly. We used the BED file to subset the BAM file, which was subsequently converted into a set of FASTA files. FASTA files contained at least 5 kbp of orthologous sequences from one or more NHP centromere assemblies. Pairs of human and NHP orthologous sequences were realigned using MAFFT (v7.453) and the following command: mafft -- maxiterate 1000 --localpair. Sequence divergence was estimated using the Tamura-Nei substitution model^94^, which accounts for recurrent mutations and differences between transversions and transitions as well as within transitions. Mutation rate per segment was estimated using Kimura’s model of neutral evolution^95^. In brief, we modeled the estimated divergence (D) is a result of between-species substitutions and within-species polymorphisms; i.e.,

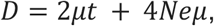

where Ne is the ancestral human effective population size, t is the divergence time for a given human– NHP pair, and μ is the mutation rate. We assumed a generation time of [20, 29] years and the following divergence times: human–macaque = [23e6, 25e6] years, human–orangutan = [12e6, 14e6] years, human–chimpanzee = [4e6, 6e6] years. To convert the genetic unit to a physical unit, our computation also assumes Ne=10,000 and uniformly drawn values for the generation and divergence times.

## DATA AVAILABILITY

The complete CHM13 chromosome 8 sequence and all CHM13 ONT data, including raw signal files (FAST5), base calls (FASTQ), and alignments (BAM/CRAM), are available at https://github.com/nanopore-wgs-consortium/chm13. In addition, the chromosome 8 sequence, CHM13 ONT FAST5 data, and CHM13 Iso-Seq data are accessioned under NCBI BioProject PRJNA559484. CHM13 PacBio HiFi data are accessioned under NCBI SRA SRX7897688, SRX7897687, SRX7897686, and SRX7897685. CHM13 Strand-seq data aligned to the CHM13 chromosome 8 assembly are accessible at doi:10.5281/zenodo.3998125. CHM13 BACs used in this study are listed in **Extended Data Table 8** with their corresponding GenBank accession numbers. Two human PacBio Iso-Seq datasets from fetal brain and testis are accessioned under NCBI BioProject PRJNA659539. The chimpanzee, orangutan, and macaque ONT FAST5 and PacBio HiFi data are accessioned under NCBI BioProject PRJNA659034.

## ACKNOWLEDGMENTS

We thank S. Goodwin (CSHL) for sequence data generation; M. Jain (UCSC) and D. Miller (UW) for re-base-calling sequence data; R. Tindell, H. Visse, A. Tornabene, and G. Ellis (UW) for technical assistance; Z. Zhao for computational assistance; F.F. Dastvan (UW) for instrumentation; D. Gordon (UW) for accessioning BACs; G. Bouffard (NHGRI) for accessioning ONT FAST5 data; J.G. Underwood (FHCRC/PacBio) for helpful discussions; and T. Brown (UW) for assistance in editing this manuscript. We acknowledge experimental support from the W. M. Keck Microscopy Center (UW) and the computational resources of the NIH HPC Biowulf cluster (https://hpc.nih.gov). This research was supported, in part, by funding from the National Institutes of Health (NIH), HG002385 and HG010169 (EEE); National Institute of General Medical Sciences (NIGMS), F32 GM134558 (GAL); Intramural Research Program of the National Human Genome Research Institute at NIH (SK, AMP, AR); National Library of Medicine Big Data Training Grant for Genomics and Neuroscience 5T32LM012419-04 (MRV); NIH/NHGRI Pathway to Independence Award K99HG011041 (PH); NIH/NHGRI R21 1R21HG010548-01 and NIH/NHGRI U01 1U01HG010971 (KHM); and the Intramural Research Program of the NIH, National Cancer Institute, Center for Cancer Research, USA (VL). EEE is an investigator of the Howard Hughes Medical Institute.

## AUTHOR CONTRIBUTIONS

GAL and EEE conceived the project; GAL, KH, KMM, AML, CB, MS generated long-read sequencing data; GAL, MRV, PH, YM, SK, SN, PCD, AR, TD, DP, AM, AVB, MK, TAG-L, CJ, SCM, KHM, and AMP analyzed sequencing data, created genome assemblies, and performed QC analyses; GAL, MRV, SK, AMP and SN finalized the chromosome 8 assembly; GAL, SK, SN, AM, AVB, and KHM assessed the assembly of the centromere; MAL generated pulsed-field gel Southern blots; GAL, LM, and MV generated microscopy data; LGD generated and analyzed droplet digital PCR data; US provided the CHM13 cell line; JGL and VL supervised experimental analyses; GAL, MRV, and EEE developed figures; and GAL and EEE drafted the manuscript.

