## Extended Data for "The structure, function, and evolution of a complete human chromosome 8"

**This PDF file includes:**

1. **Extended Data Figures 1 to 16**
2. **Extended Data Tables 1 to 8**
3. **References**

**EXTENDED DATA FIGURES**

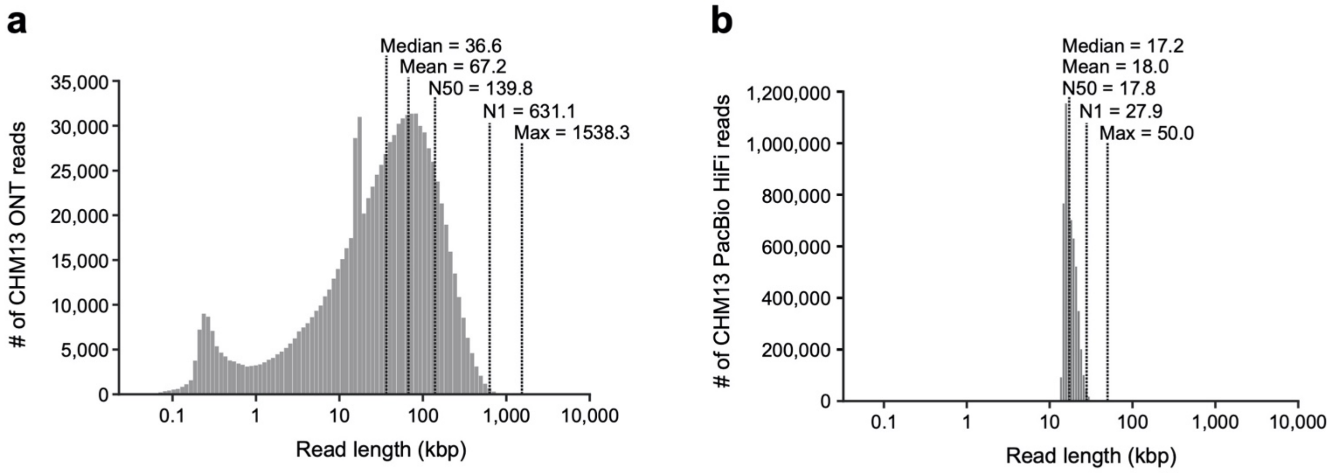


**Extended Data Figure 1. Ultra-long ONT and PacBio HiFi data from the CHM13 genome.**
**a,b)** Read-length distributions of **a)** ultra-long ONT and **b)** PacBio HiFi data generated from the CHM13 genome for this study. The median, mean, N50, N1, and max read lengths are indicated.

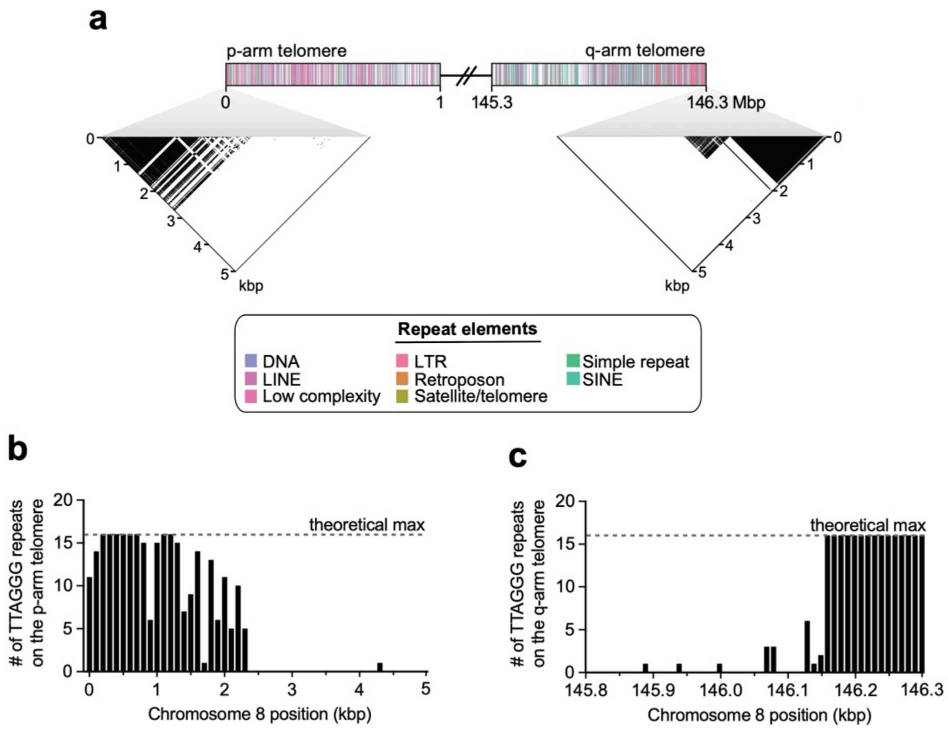


**Extended Data Figure 2. Chromosome 8 telomeres. a)** Schematic showing the first and last megabase of the CHM13 chromosome 8 assembly. A dot plot of the terminal 5 kbp shows high sequence identity among the last ~2.5 kbp of the chromosome, consistent with the presence of a high-identity telomeric repeating unit. **b,c)** Frequency of the TTAGGG telomeric repeat in the last 5 kbp of the p- (**Panel a**) and q- (**Panel b**) arms in chromosome 8. The p-arm has a gradual transition to pure TTAGGG repeats over nearly 1 kbp, while the q-arm has a very sharp transition to pure TTAGGG repeats that occurs over nearly 300 bp.

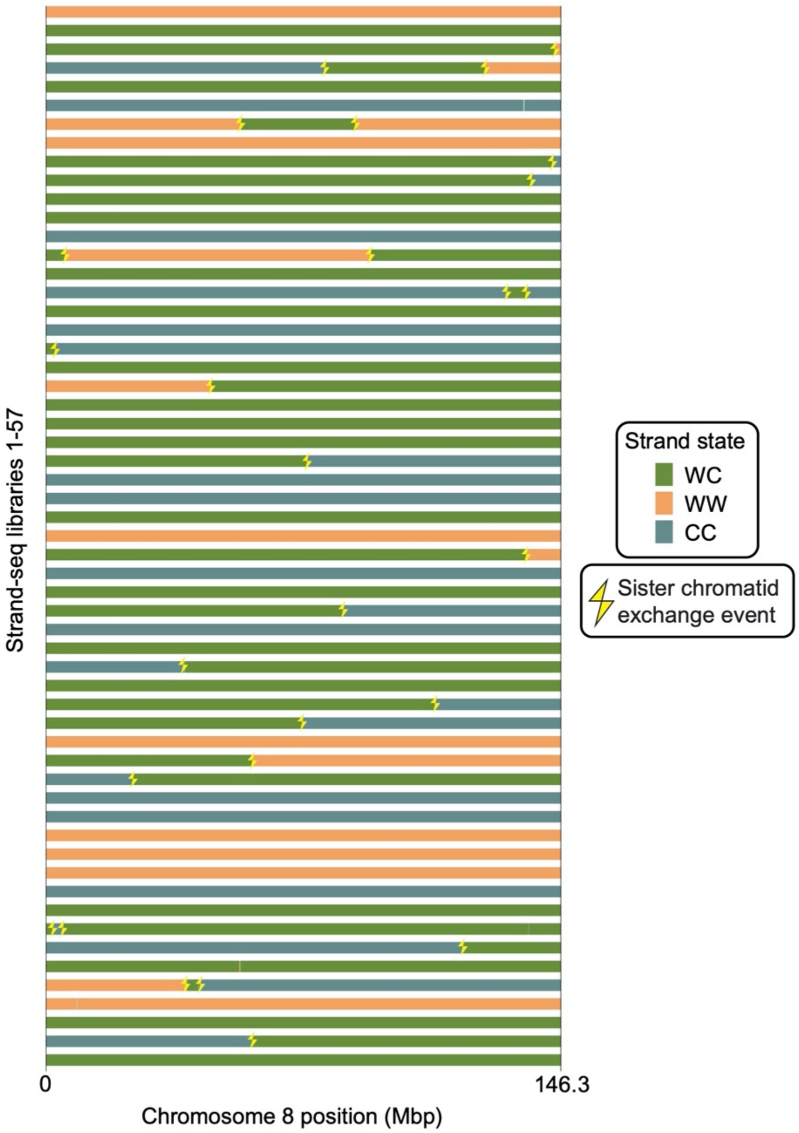


**Extended Data Figure 3. Strand-seq validation of the chromosome 8 assembly.** Strand-seq is a single-cell sequencing technique able to assess directional and structural contiguity of individual homologs by sequencing only template single-stranded DNA[1–3]. Horizontal colored bars show Strand-seq strand states for 57 libraries. Such strand state changes are normally caused by a double-strand-break that occurred during DNA replication and was repaired by a sister chromatid[3]. Observed strand state changes are randomly distributed along each Strand-seq library (yellow thunderbolts) for chromosome 8 and, thus, are not indicative of a genome misassembly. Genome misassemblies are indicated by a recurrent change in strand state over the same region in multiple Strand-seq cells[4]. WC: Watson-Crick; WW: Watson-Watson; CC: Crick-Crick.

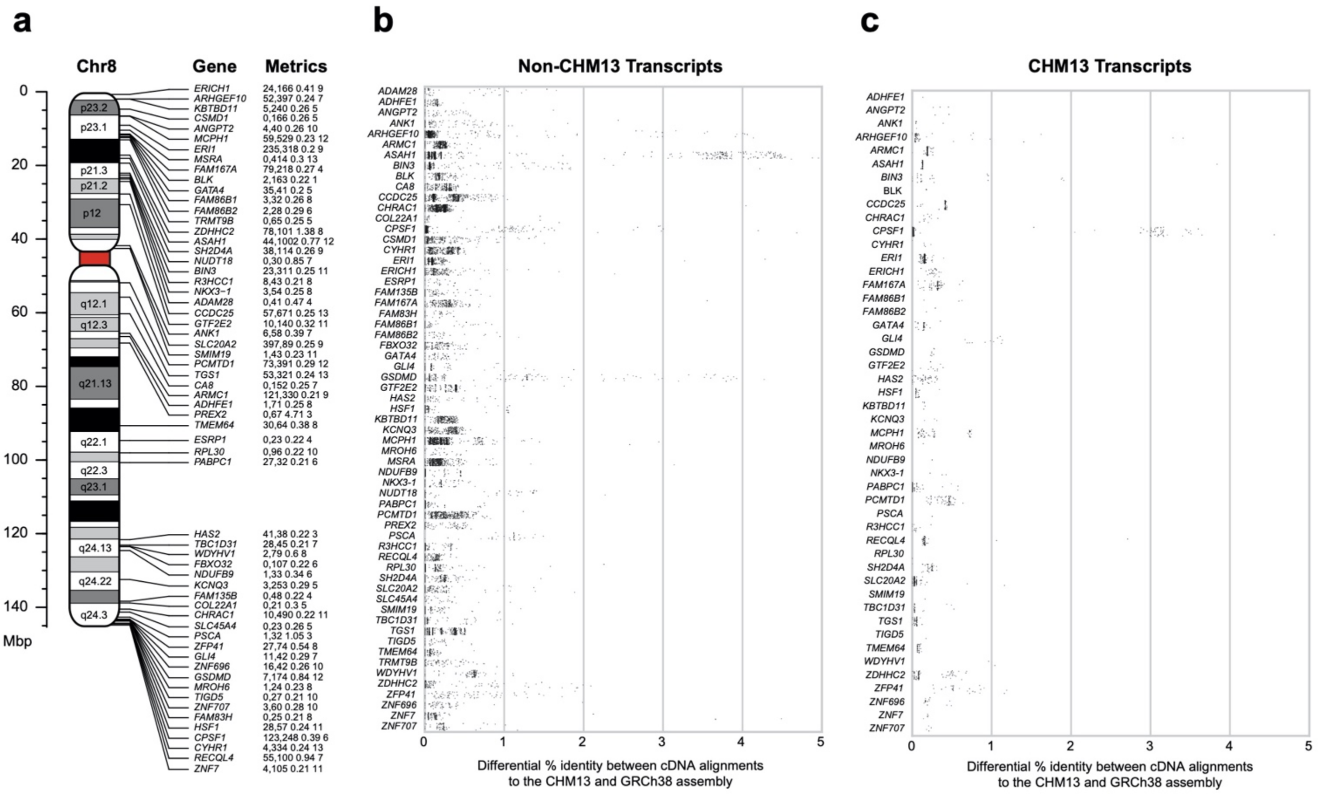


**Extended Data Figure 4. Protein-coding genes with improved alignment to the CHM13 chromosome 8 assembly relative to GRCh38. a)** Ideogram of chromosome 8 showing protein-coding genes with improved transcript alignments to the CHM13 chromosome 8 assembly relative to GRCh38 (hg38). Each gene is labeled with its name, count of improved transcripts from CHM13 cell line, other tissues, the average percent improvement of non-CHM13 cell line alignments, and the number of tissue sources with improved transcript mappings. **b,c)** Differential percent identity of cDNA alignments to CHM13 chromosome 8 compared to GRCh38 for **b)** non-CHM13 cell line transcripts and **c)** CHM13 cell line transcripts. See **Extended Data Fig. 16** for examples of protein-coding genes with higher sequence identity to the CHM13 chromosome 8 assembly than to GRCh38.

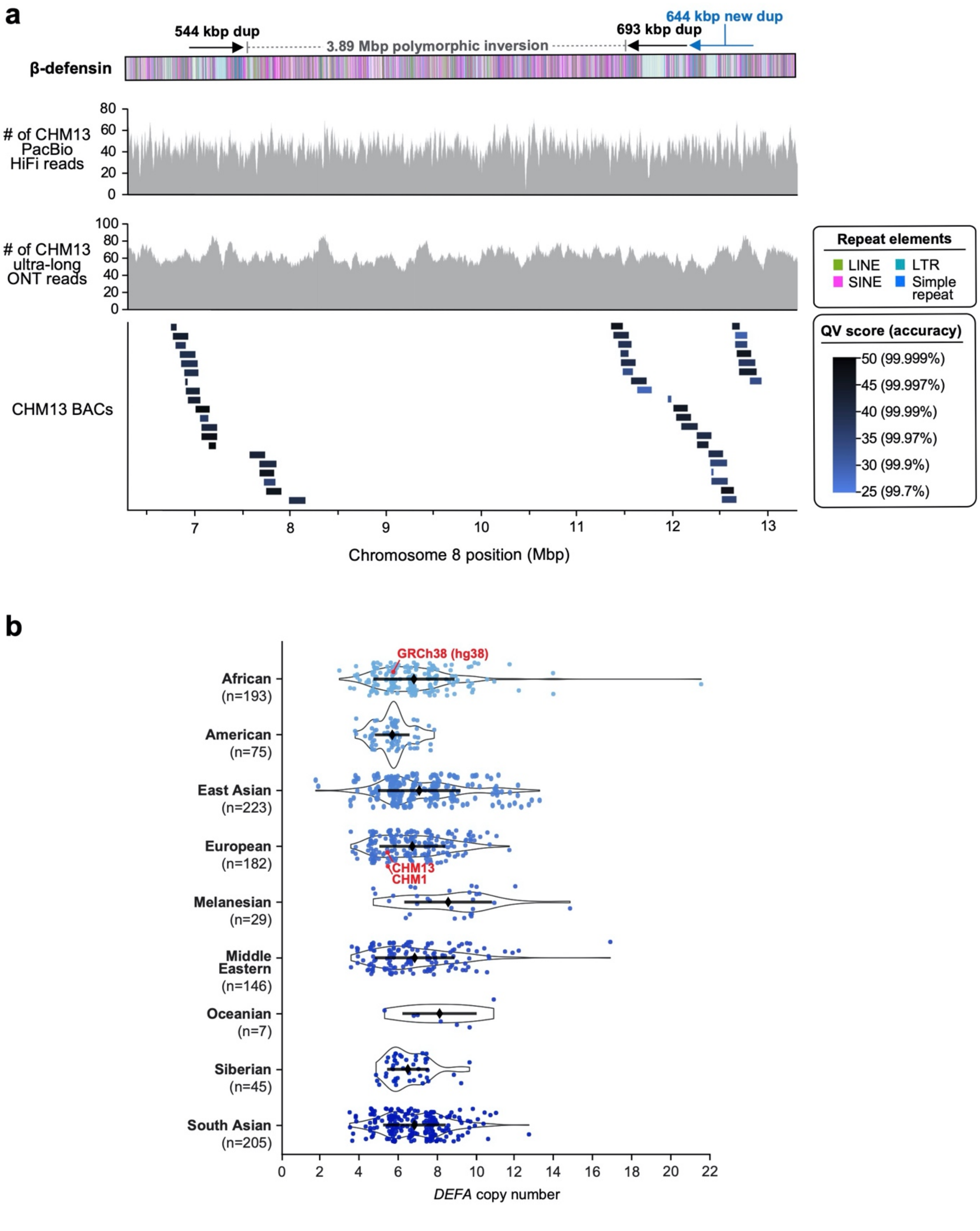


**Extended Data Figure 5. Validation of the CHM13 β-defensin locus, and copy number of the *DEFA* gene family. a)** Coverage of CHM13 ONT and PacBio HiFi data along the CHM13 β-defensin locus (top two panels). The ONT and PacBio data have largely uniform coverage, indicating it is free of large structural errors. The dip in HiFi coverage near position 10.46 Mbp is due to a G/A bias in HiFi chemistry[5]. The alignment of 47 CHM13 BACs (bottom panel) reveals that those regions have an estimated QV score >25 (>99.7% accurate). **b)** Copy number of *DEFA* [chr8:6976264−6995380 in GRCh38 (hg38)] throughout the human population.

**a**

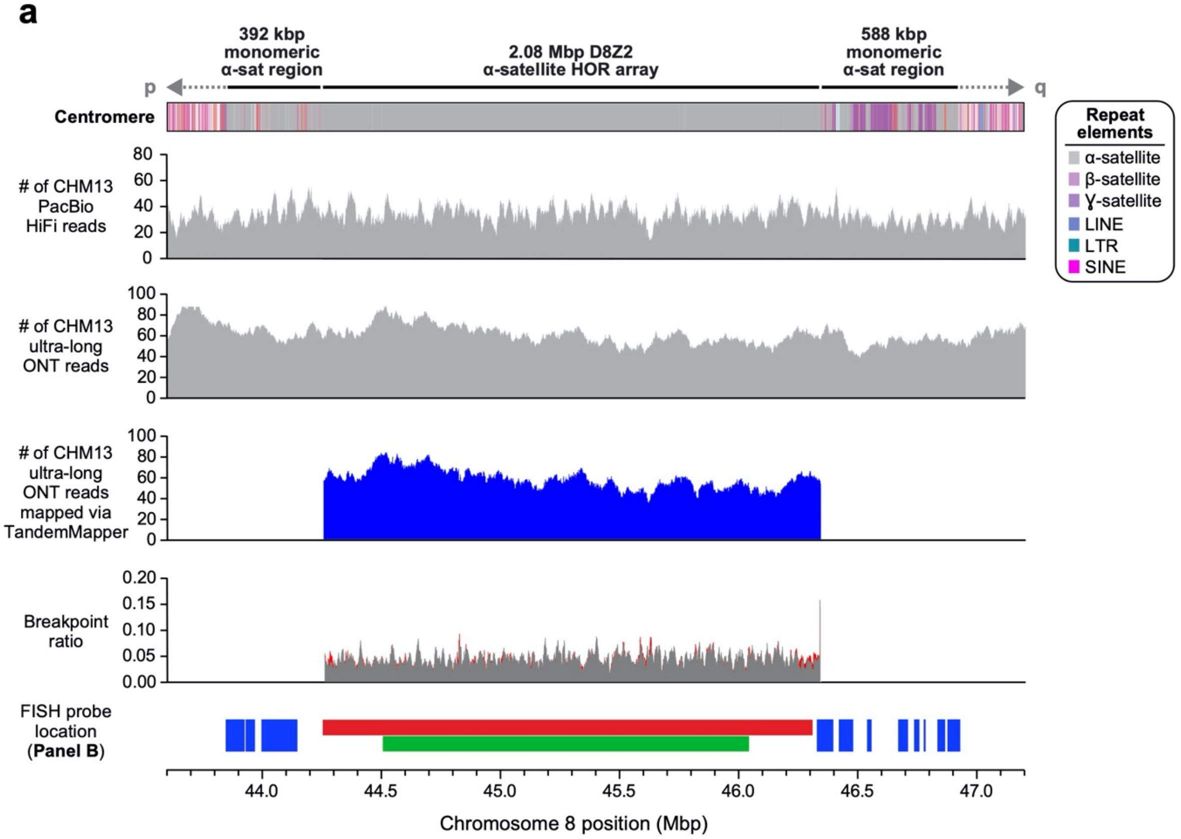


**b**

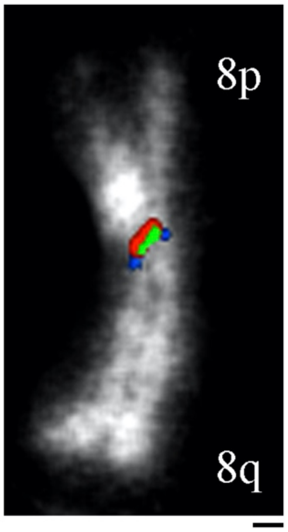


**c**

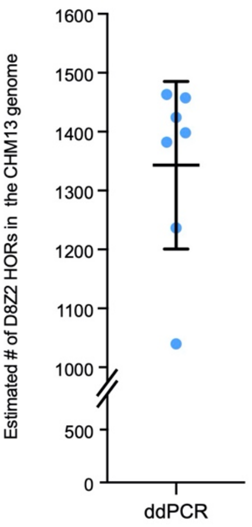


**Extended Data Figure 6. Validation of the CHM13 centromeric region. a)** Coverage of CHM13 ONT and PacBio HiFi data along the CHM13 centromeric region (top two panels) is largely uniform, indicating a lack of large structural errors. Analysis with TandemMapper and TandemQUAST[6], which are tools that assess repeat structure via mapped reads (third panel) and misassembly breakpoints (fourth panel; red), indicates that the chromosome 8 D8Z2 HOR array lacks large-scale assembly errors. Three different FISH probes targeting regions in the chromosome 8 centromeric region (bottom panel) are used to confirm the organization of the α-satellite DNA (**Panel b**). **b)** Representative FISH image of a stretched CHM13 chromosome showing the predicted organization of the chromosome 8 centromeric region (**Panel a**). **c)** Droplet digital PCR (ddPCR) of the chromosome 8 D8Z2 array indicates that there are 1344 +/- 142 D8Z2 HORs present on chromosome 8, consistent with the predictions from an *in silico* restriction digest and StringDecomposer analysis (**Methods**). Mean +/- s.d. is shown. Bar = 5 microns.

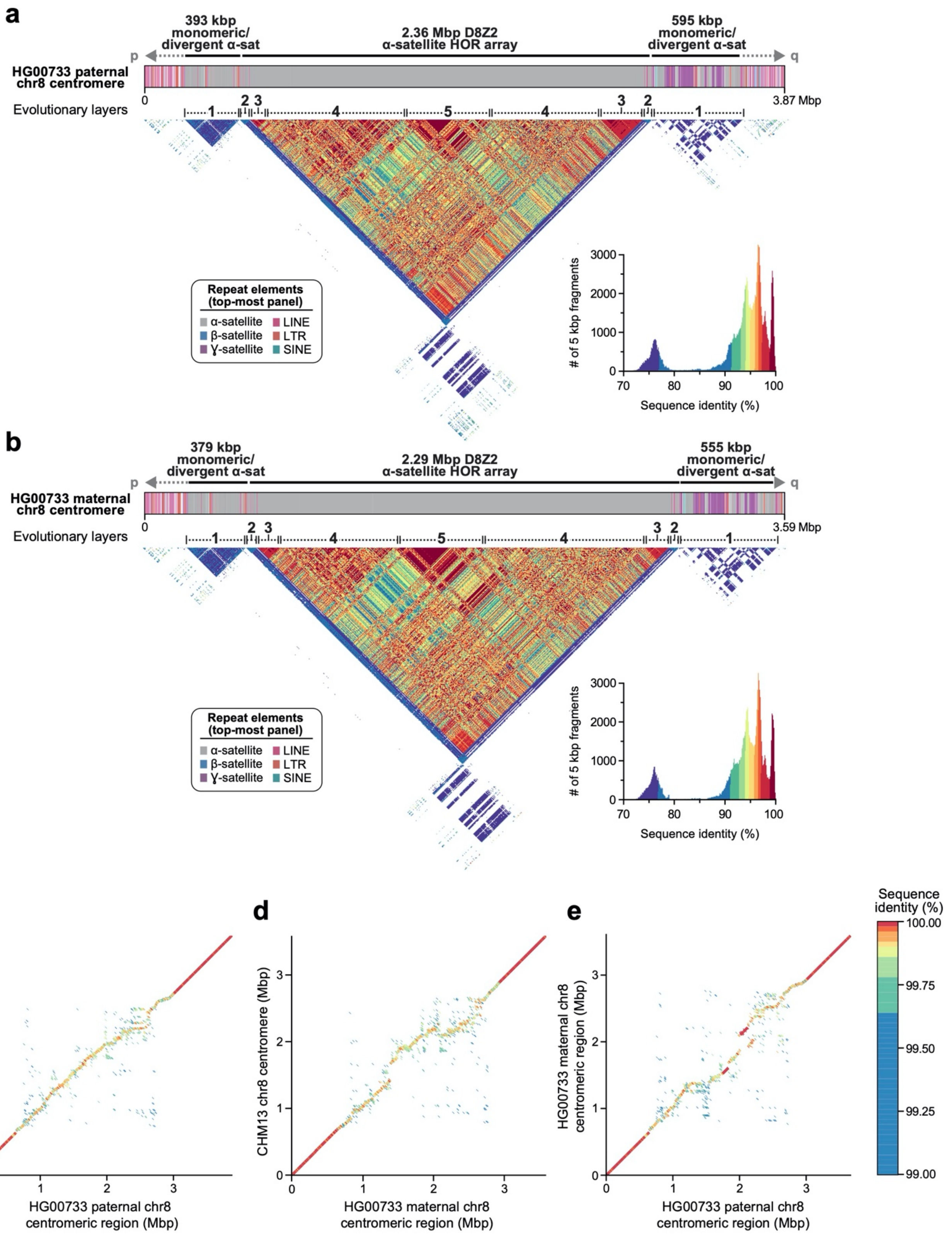


**Extended Data Figure 7. Assembly and comparison of the chromosome 8 centromeric regions from a Puerto Rican child (HG00733) to the CHM13 centromeric region. a,b)** Repeat structure and sequence identity heat map of the **a)** paternal and **b)** maternal chromosome 8 centromeric regions from a diploid genome (HG00733) shows structural and evolutionary similarity to the CHM13 chromosome 8 centromeric region (**Fig. 2a**). **c-e)** Dot plot comparisons between the **c)** CHM13 and paternal, **d)** CHM13 and maternal, and **e)** paternal and maternal chromosome 8 centromeric regions in a diploid genome (HG00733) shows >99% sequence identity overall, with high concordance in the unique and monomeric α-satellite regions of the centromeres (dark red line) that devolves into lower sequence identity in the α-satellite HOR array, consistent with rapid evolution of this region.

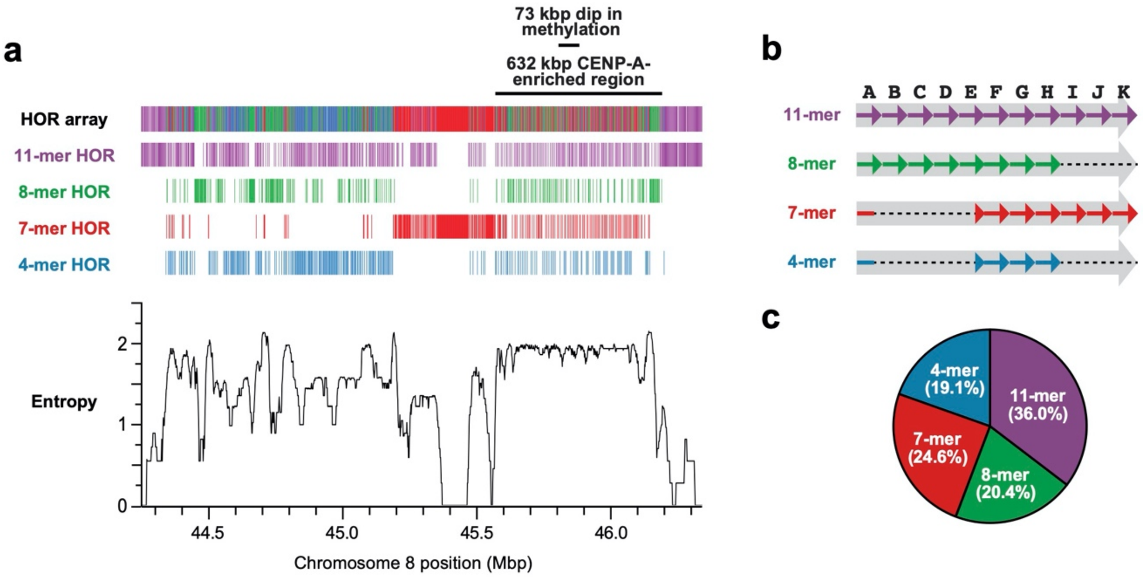


**Extended Data Figure 8. Composition, organization, and entropy of the CHM13 D8Z2 α-satellite HOR array. a)** HOR composition and organization of the chromosome 8 α-satellite array as determined via StringDecomposer[7]. The predominant HOR subtypes (4-, 7-, 8-, and 11-mers) are shown, while those occurring less than 15 times are not (see **Methods** for absolute quantification). The entropy of the D8Z2 HOR array is plotted in the bottom panel and reveals that the hypomethylated and CENP-A-enriched regions have the highest consistent entropy in the entire array. **b)** Organization of α-satellite monomers within each HOR. The initial monomer of the 4- and 7-mer HORs is a hybrid of the A and E monomers, with the first 87 bp the A monomer and the subsequent 84 bp the E monomer.
**c)** Abundance of the predominant HOR types within the D8Z2 HOR array as determined via StringDecomposer[7].

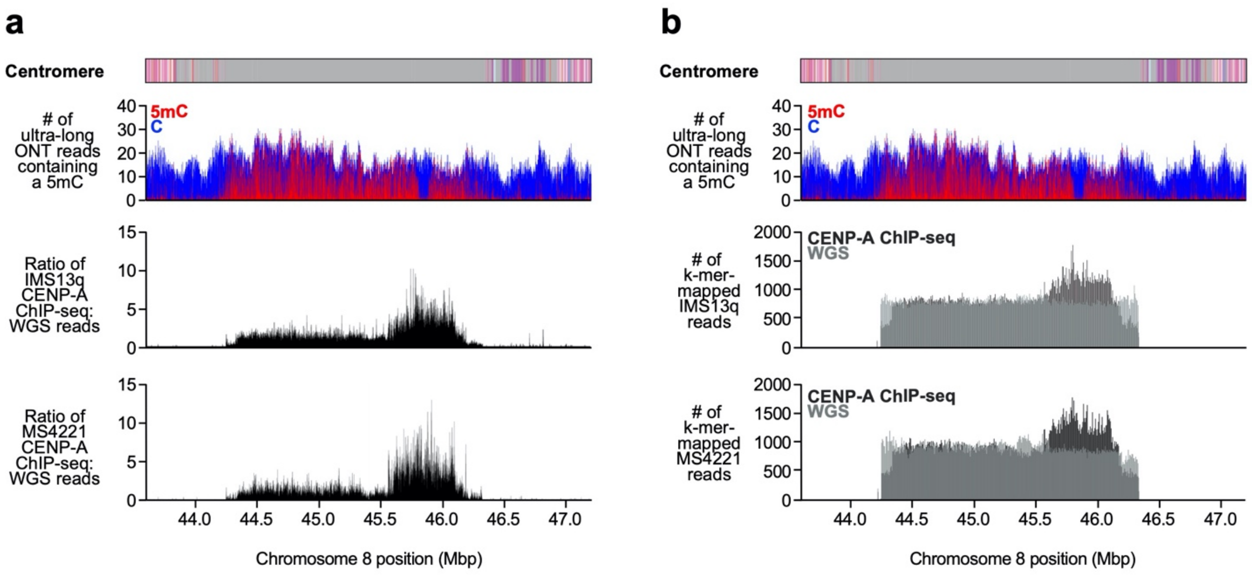


**Extended Data Figure 9. Location of CENP-A chromatin within the CHM13 D8Z2 HOR array.** **a,b)** Plot of **a)** the ratio CENP-A ChIP-seq reads to WGS reads mapped via BWA-MEM, or **b)** the number of k-mer-mapped CENP-A ChIP-seq (black) or WGS (dark gray) reads from diploid neocentromeric human cell lines (**Methods**). While these cell lines have a neocentromere located on either chromosome 13 (IMS13q) or 8 (MS4221)[8], they both have at least one karyotypically normal chromosome 8 from which centromeric chromatin can be mapped. We limited our analysis to diploid human cell lines rather than aneuploid ones to avoid potentially confounding results stemming from multiple chromosome 8 copies that vary in structure, such as those observed in HeLa cells[9].

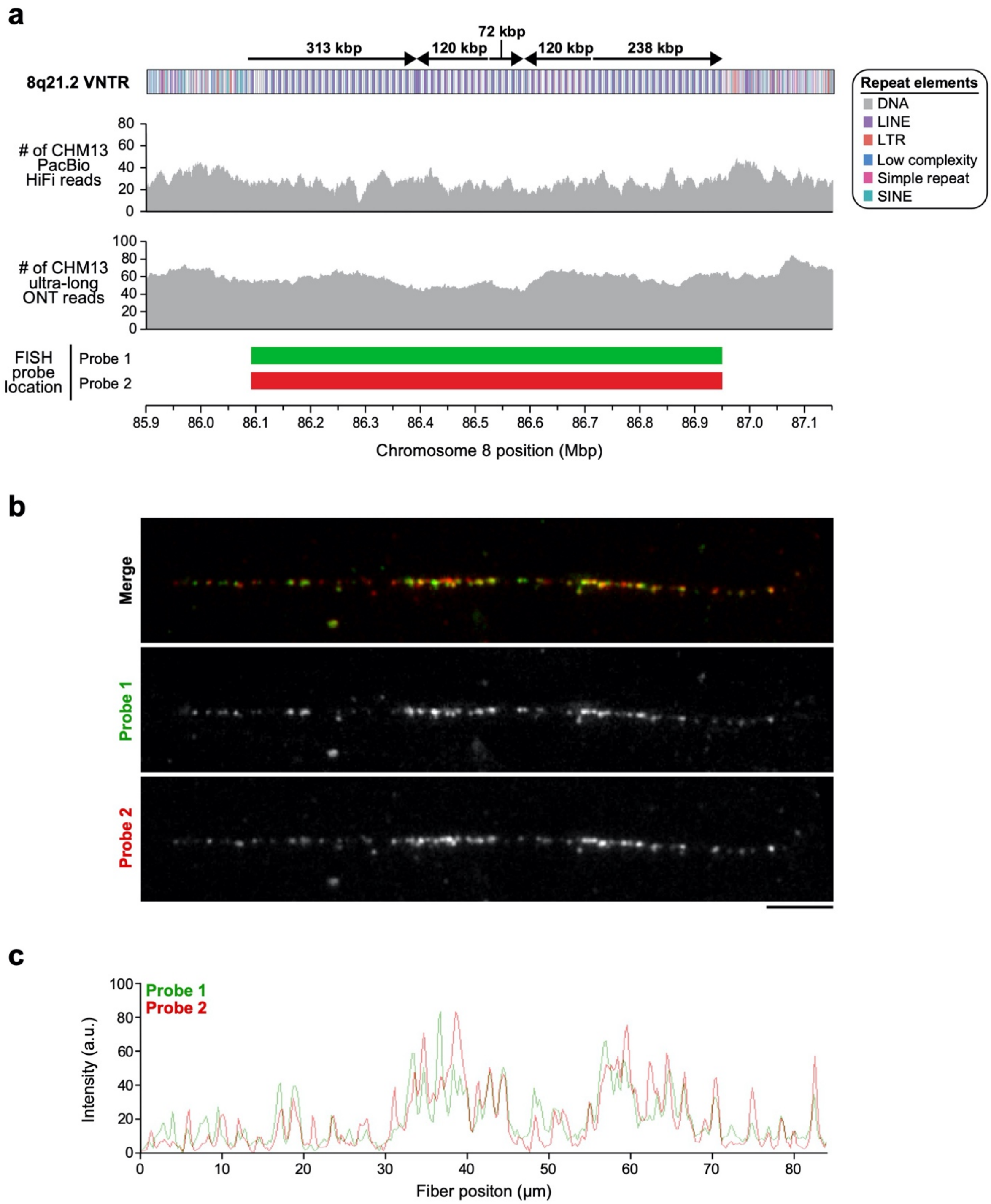


**Extended Data Figure 10. Validation of the CHM13 8q21.2 VNTR. a)** Coverage of CHM13 ONT and PacBio HiFi data along the 8q21.2 VNTR (top two panels) is largely uniform, indicating a lack of large structural errors. Two FISH probes targeting the 12.192 kbp repeat in the 8q21.2 VNTR are used to estimate the number of repeats in the CHM13 genome (**Panels b,c**). **b)** Representative FISH images of a CHM13 stretched chromatin fiber. Although the FISH probes were designed against the entire VNTR array, stringent washing during FISH produces a punctate probe signal pattern, which may be due to stronger hybridization of the probe to a specific region in the 12.192 kbp repeat (perhaps based on GC content or a lack of secondary structures). This punctate pattern can be used to estimate the repeat copy number in the VNTR, thereby serving as a source of validation. **c)** Plot of the signal intensity on the CHM13 chromatin fiber shown in **Panel b**. Quantification of peaks across three independent experiments reveals an average of 63 +/- 7.55 peaks and 67 +/- 5.20 peaks from the green and red probes, respectively, which is consistent with the number of repeat units in the 8q21.2 assembly (67 full and 7 partial repeats). Bar = 5 micron.

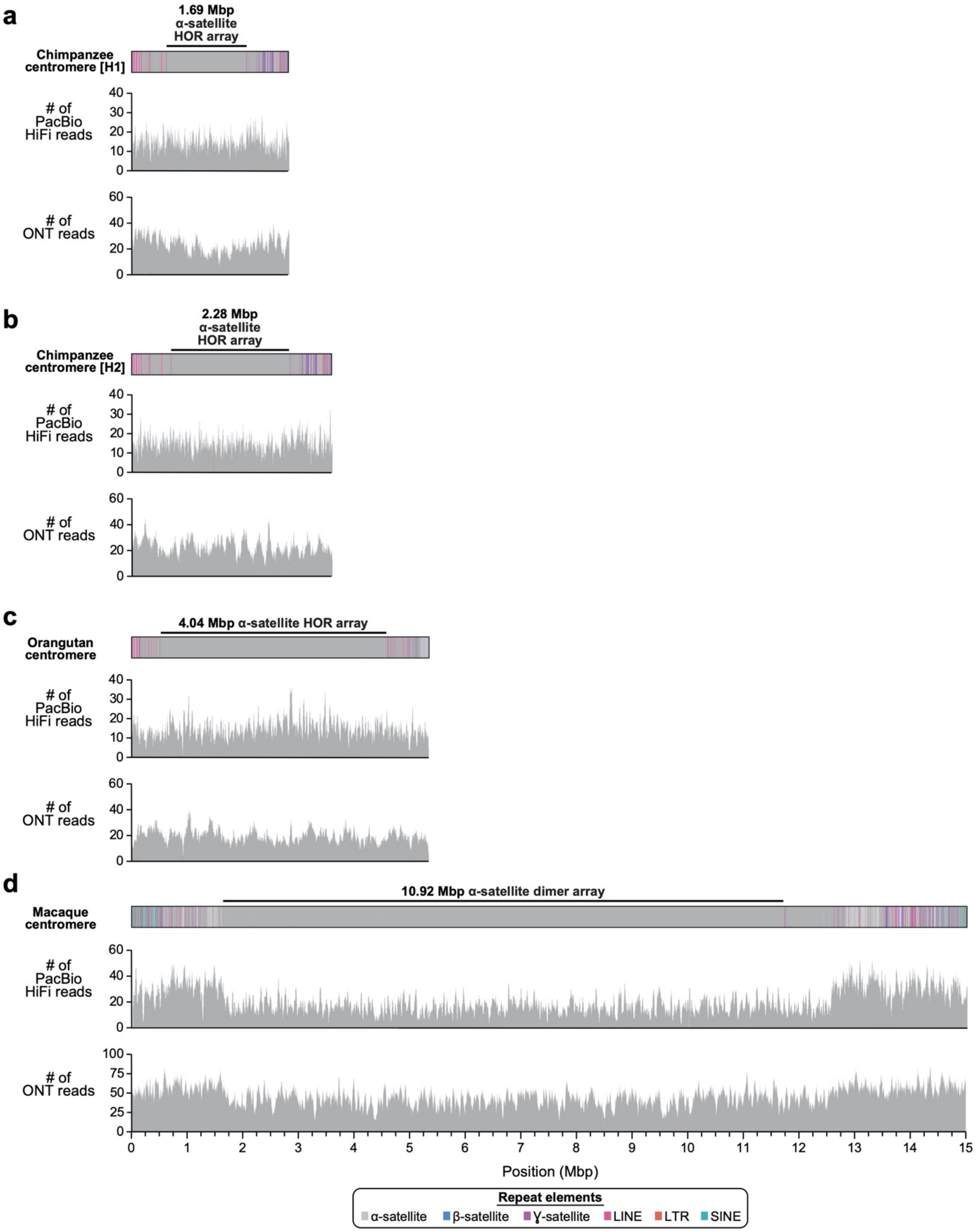


**Extended Data Figure 11. Validation of the NHP centromeric regions with mapped long reads. a-d)** Coverage of the **a)** chimpanzee (H1), **b)** chimpanzee (H2), **c)** orangutan, and **d)** macaque chromosome 8 centromeric regions with PacBio HiFi and ONT data generated from the same genome reveals largely uniform coverage. The increase in coverage on the edges of the macaque centromeric region is due to the presence of only one haplotype representation in these regions. All assemblies are to scale. H1, haplotype 1; H2, haplotype 2.

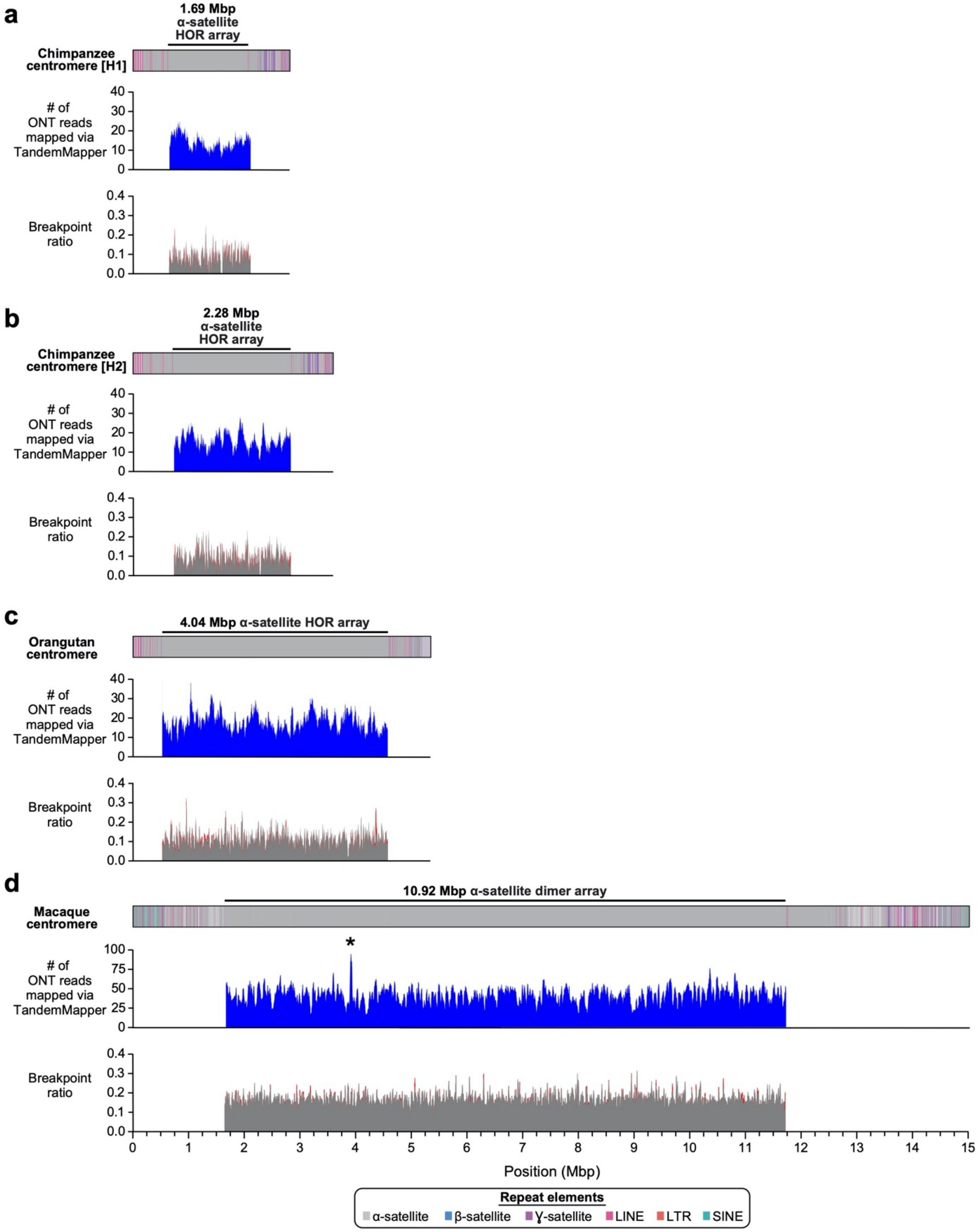


**Extended Data Figure 12. Validation of the NHP centromeric regions with TandemQUAST analysis. a-d)** ONT reads mapped with TandemMapper (top panel) and breakpoint ratios identified with TandemQUAST (bottom panel) for **a)** chimpanzee (H1), **b)** chimpanzee (H2), **c)** orangutan, and **d)** macaque chromosome 8 centromeric regions reveals a lack of large structural errors, except for a potential collapse identified in the macaque centromere (marked with an asterisk). All assemblies are to scale. H1, haplotype 1; H2, haplotype 2.

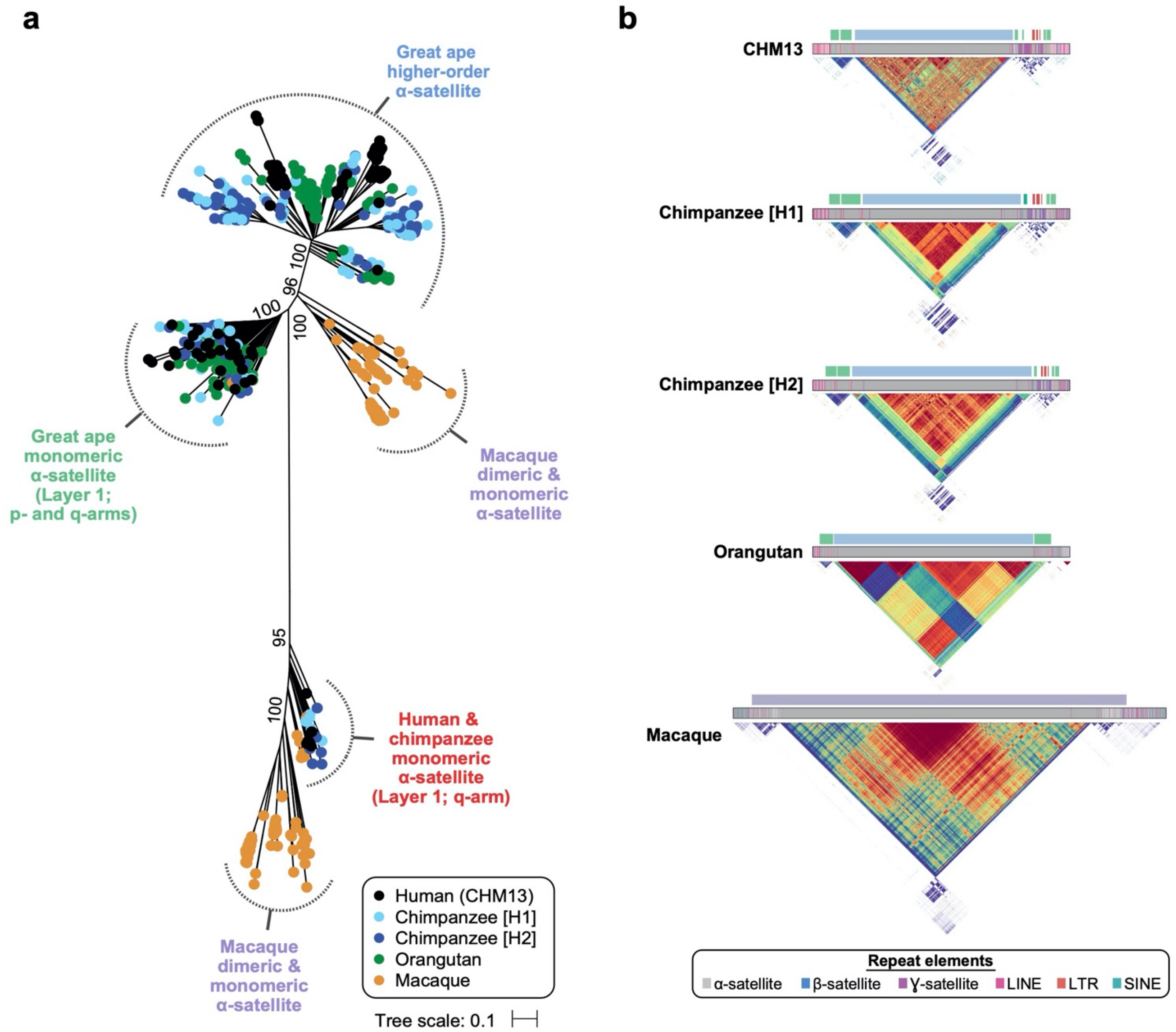


**Extended Data Figure 13. Relative position of α-satellite phylogenetic clades. a)** Phylogenetic tree with bootstrapping indicated at each major node. The clades are color-coded, and the relative position of the α-satellite in each clade is indicated in **Panel b**. **b)** Location of α-satellite from each clade from **Panel a**. The ancient monomeric α-satellite (red) is restricted to the q-arm in CHM13 and chimpanzee and represents the vestigial centromeric α-satellite from the common ancestor with macaque.

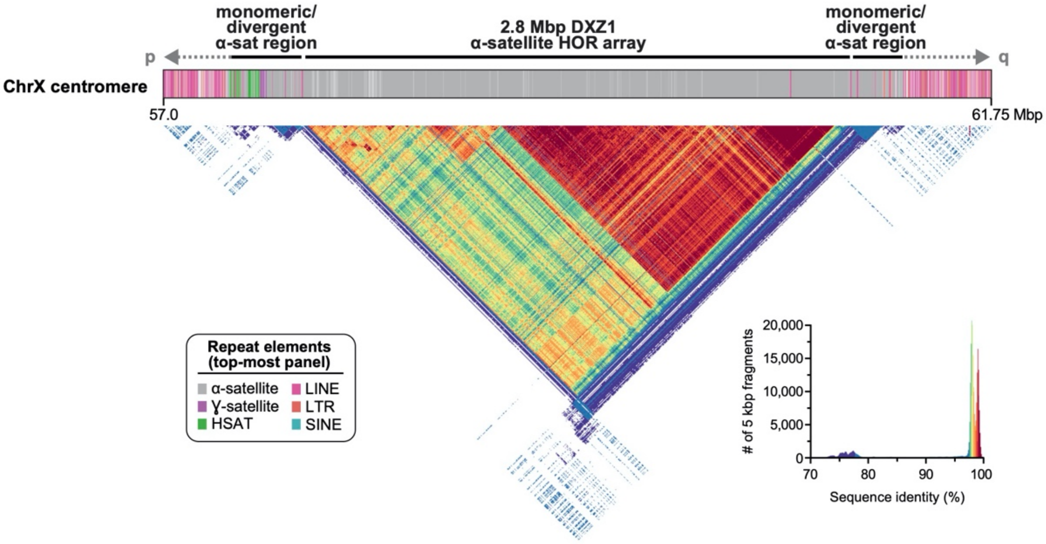


**Extended Data Figure 14. Sequence composition and identity of the chromosome X centromeric region.** Repeat structure and sequence identity heat map of the chromosome X centromeric region reveals >90% sequence identity across the HOR array and four distinct evolutionary layers that lack the symmetricity observed at the chromosome 8 HOR array, in concordance with prior analysis[10].

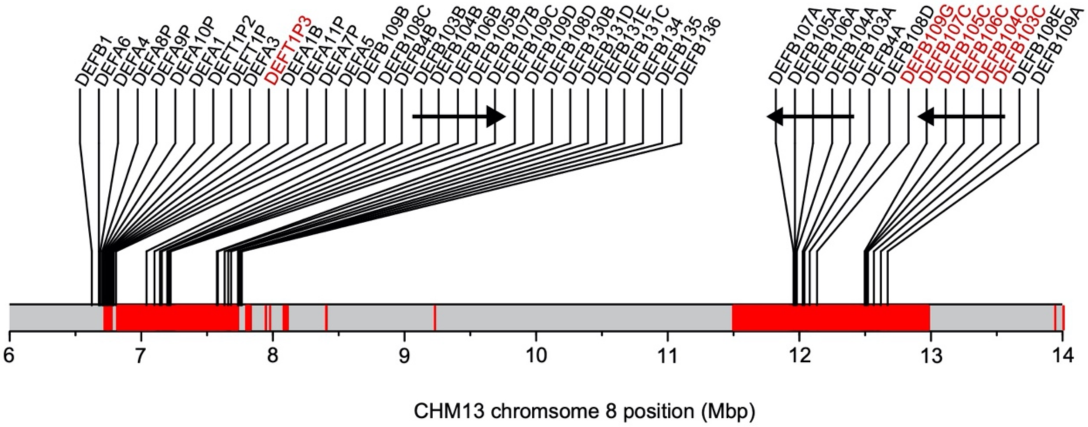


**Extended Data Figure 15. Location of *DEFA* and *DEFB* genes in the CHM13 chromosome 8 β-defensin locus.** SD regions as identified by SEDEF[11], and new paralogs are shown in red. Duplication cassettes are marked with arrows indicating orientation for each copy.

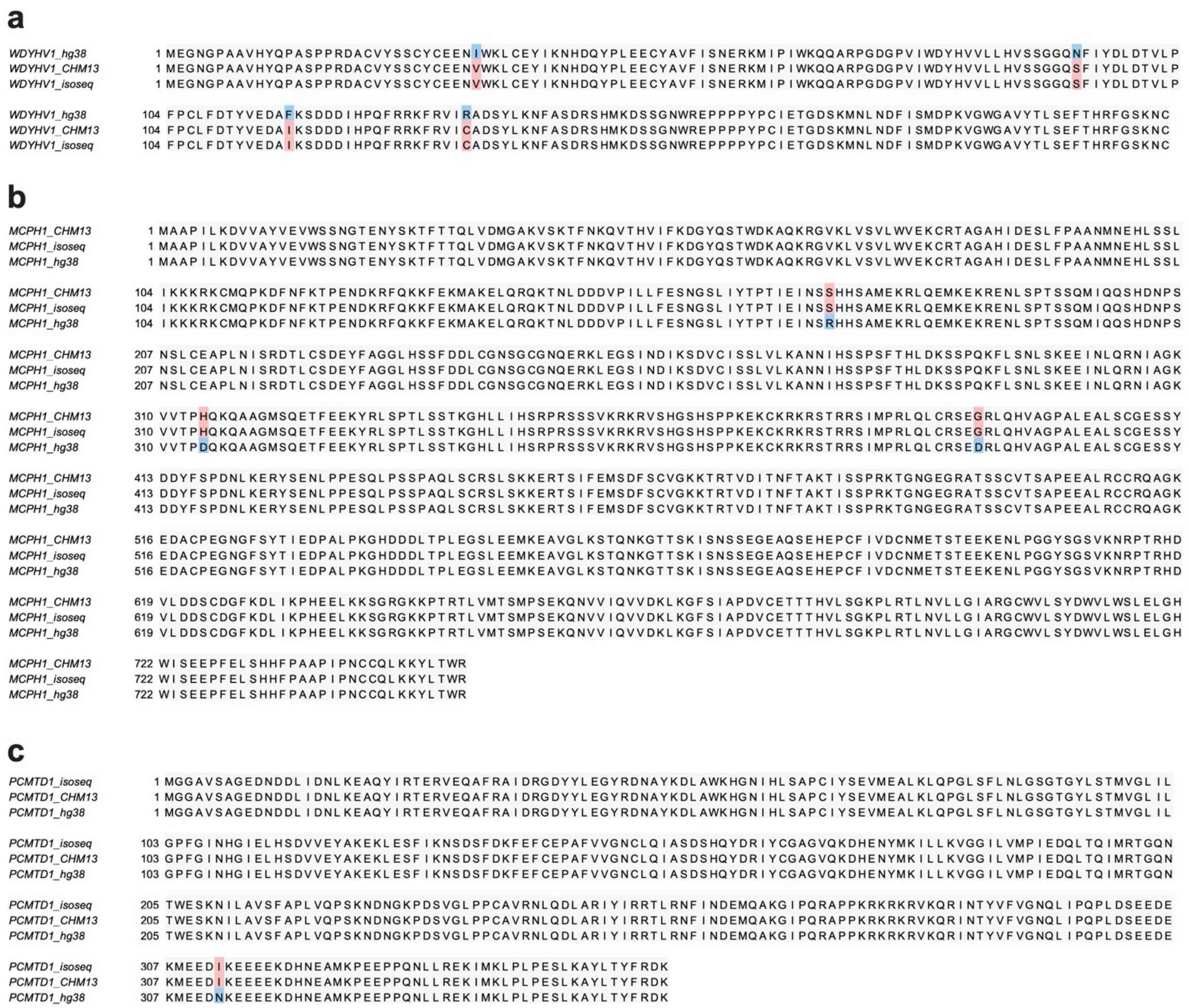


**Extended Data Figure 16. Multiple-sequence alignments of predicted open reading frames (ORFs) for genes that align better to the CHM13 chromosome 8 relative to GRCh38. a-c)** Multiple-sequence alignments for **a)** *WDYHV1*, **b)** *MCPH1*, and **c)** *PCMTD1*, all of which have at least 0.1% greater sequence identity of >20 full-length Iso-Seq transcripts to the CHM13 chromosome 8 assembly than to GRCh38 (**Methods**). For each gene, the GRCh38 annotation is compared to the same annotation lifted over to the CHM13 chromosome 8 assembly, and the substitutions are confirmed by translated predicted ORFs from Iso-Seq transcripts. Matching amino acids are shaded in gray, those matching only the Iso-Seq data are in red, and those different from the Iso-Seq data are in blue. Each substitution in CHM13 relative to GRCh38 has an allele frequency of 0.36 in gnomAD (v3).

### EXTENDED DATA TABLES

**Extended Data Table 1. Differences in CHM13 and GRCh38 (hg38) chromosome 8 *DEFA* and *DEFB* genes.** See accompanying Excel file.

**Extended Data Table 2. Evolutionary layers within the CHM13 and Puerto Rican child chromosome 8 centromeres.**

| Evolutionary layer | CHM13 chromosome 8 centromere (bp) | | | HG00733 paternal chromosome 8 centromere (bp) | | | HG00733 maternal chromosome 8 centromere (bp) | | |
|---|---|---|---|---|---|---|---|---|---|
| | p-arm | q-arm | Total | p-arm | q-arm | Total | p-arm | q-arm | Total |
| 1 | 323918 | 496831 | 820749 | 319969 | 496835 | 816804 | 319966 | 496808 | 816774 |
| 2 | 59301 | 57889 | 117190 | 59306 | 57925 | 117231 | 59301 | 57925 | 117226 |
| 3 | 92405 | 149484 | 241889 | 94781 | 261407 | 356188 | 124624 | 149808 | 274432 |
| 4 | 842106 | 577929 | 1420035 | 816758 | 780819 | 1597577 | 630283 | 971332 | 1601615 |
| 5 | -- | -- | 416216 | -- | -- | 408974 | -- | -- | 417771 |

**Extended Data Table 3. Sequence and assembly of NHP genomes.**

| Species | Assembly* | | | PacBio HiFi data | | ONT data | |
|---|---|---|---|---|---|---|---|
| | Size (Gbp) | No. of contigs | N50 (Mbp) | Sequencing depth† | Read N50 (kbp) | Sequencing depth† | Read N50 (kbp) |
| Chimpanzee (*Pan troglodytes*; Clint; S006007) | 6.02 | 26,305 | 57.99 | 40.08 | 11 | 48.04 | 67 |
| Orangutan (*Pongo abelii*; Susie; PR01109) | 6.02 | 10,890 | 3.6 | 24.71 | 17.9 | 39.69 | 63.2 |
| Macaque (*Macaca mulatta*; AG07107) | 6.12 | 19,526 | 1.9 | 27.91 | 19.2 | 56.5 | 33.3 |

*Assembled from PacBio HiFi data with HiCanu (Nurk et al., Genome Res, 2020)

†Assumes a 3.2 Gbp genome for each species

**Extended Data Table 4. Chromosome 8 centromeric mutation rate.** See accompanying Excel file.

**Extended Data Table 5. Heterozygous sites within human chromosome 8.**

| CHM13 chromosome 8 coordinate | Insertion size (bp) | % of ONT reads >50 kbp long supporting the CHM13 chromosome 8 assembly | % of ONT reads >50 kbp long supporting the alternate structure |
|---|---|---|---|
| chr8:21,025,201 | 8829-8923 | 58.33% (35/60) | 41.67% (25/60) |
| chr8:80,044,843 | 7884 | 98.59% (70/71) | 1.41% (1/71) |
| chr8:121,388,618 | 5928-6023 | 70% (42/60) | 30% (18/60) |

ONT: Oxford Nanopore Technologies

**Extended Data Table 6. PacBio Iso-Seq datasets.** See accompanying Excel file.

**Extended Data Table 7. Genes with greater sequence identity to CHM13 chromosome 8 than GRCh38.** See accompanying Excel file.

**Extended Data Table 8. CHM13 BACs used in this study.** See accompanying Excel file.

### EXTENDED DATA REFERENCES
